# A validated antibody toolbox for ALS research

**DOI:** 10.1101/2025.09.14.676084

**Authors:** Riham Ayoubi, Emma J. MacDougall, Ian McDowell, Michael S. Biddle, Bárbara T. Ferreira, CongYao Zha, Marie-France Dorion, Jay P. Ross, Sara González Bolívar, Vera Ruiz Moleón, Charles Alende, Vincent Francis, Maryam Fotouhi, Mathilde Chaineau, Carol X.-Q. Chen, Valerio E.C. Piscopo, Vincent Soubannier, Tracy Keates, Wen Hwa Lee, Brian D. Marsden, Leonidas Koukouflis, Edvard Wigren, Carolyn A. Marks, Luke M. Healy, Patrick A. Dion, Guy A. Rouleau, Edward A. Fon, Harvinder S. Virk, Susanne Gräslund, Opher Gileadi, Aled M. Edwards, Thomas M. Durcan, Peter S. McPherson, Carl Laflamme

## Abstract

A substantial fraction of amyotrophic lateral sclerosis (ALS)-associated proteins remain poorly characterized, in part because of the limited availability of validated research antibodies. We established knockout (KO)-based antibody characterization workflows and demonstrated that widely used antibodies against the major ALS-associated protein C9orf72 lacked specificity (Laflamme et al., 2019). We subsequently scaled this framework to systematically benchmark research antibodies, revealing that up to 61% fail to perform as recommended by manufacturers (Ayoubi et al., 2023). Here, we extend this approach by establishing the ALS-Reproducible Antibody Platform (ALS-RAP), a comprehensive effort to generate a publicly available dataset of KO-validated antibodies targeting proteins encoded by ALS risk genes. In total, we characterized 303 antibodies against 33 ALS-associated proteins to identify high-quality reagents for use in western blot, immunoprecipitation, and immunofluorescence. Using these antibodies, we profiled protein levels across human induced pluripotent stem cell (iPSC)-derived and primary neurological cell types. These analyses revealed diverse cellular distributions and higher levels of several ALS-associated proteins in glial populations, consistent with emerging evidence for immune contributions to ALS. Together, ALS-RAP provides a validated antibody toolbox and protein expression resource to support the study of ALS-associated proteins.

## Introduction

Research antibodies are among the most widely used tools for protein detection and quantification in biomedical research, yet their performance and specificity are often insufficiently characterized [1–4]. Building on our previous studies establishing knockout (KO)-based antibody characterization workflows and benchmarking antibody performance [5, 6], we applied this framework to proteins encoded by amyotrophic lateral sclerosis (ALS) risk genes, a genetically defined but mechanistically heterogeneous disease. More than 30 genes have been linked to familial and sporadic ALS [7, 8], yet most ALS-associated proteins remain poorly characterized at the protein level. This gap reflects both the limited availability of validated reagents and the lack of systematic protein-level characterization across relevant neurological cell types.

To address this limitation, we established the ALS-Reproducible Antibody Platform (ALS-RAP), a systematic effort to characterize antibodies targeting ALS-associated proteins using KO-based workflows [9], with a focus on renewable reagents (monoclonal and recombinant antibodies). ALS-RAP builds on antibody characterization strategies developed through the Antibody Characterization through Open Science (YCharOS) initiative, an open-science ecosystem that brings together academic laboratories and antibody manufacturers to systematically evaluate research antibodies and openly disseminate characterization datasets [10–12]. To expand renewable reagent coverage, the Structural Genomics Consortium (SGC) generated recombinant antibodies for several ALS targets lacking suitable commercial reagents [13].

Through ALS-RAP, we assembled and characterized antibodies targeting 33 ALS-associated proteins and identified high-quality renewable reagents suitable for common applications. These validated antibodies enabled us to examine protein levels across human induced pluripotent stem cell (iPSC)-derived and primary neurological cell types. Together, this work extends antibody characterization to a disease-focused protein landscape and provides a resource for investigating these proteins in relevant cellular contexts.

## Results

### Genetic prioritization and initial characterization of ALS-RAP targets

We prioritized 33 ALS-related genes supported by replicated genetic evidence: *ACSL5*, *ALS2*, *ANG*, *ANXA11*, *ATXN2*, *C9orf72*, *CAV1*, *CCNF*, *CHCHD10*, *CHMP2B*, *FIG4*, *FUS*, *HNRNPA1*, *HNRNPA2B1*, *KIF5A*, *LGALS1*, *MATR3*, *NEK1*, *OPTN*, *PFN1*, *SETX*, *SIGMAR1*, *SOD1*, *SPG11*, *SQSTM1*, *TAF15*, *TARDBP*, *TBK1*, *TIA1*, *TUBA4A*, *UBQLN2*, *VAPB*, *VCP*. Building on a previous classification framework [7], these genes were grouped according to variant frequency and estimated pathogenic impact, including rare variants with high, intermediate, or small effect sizes, as well as rare or common variants with uncertain impact. Of these, 25 genes have also been curated by the Clinical Genome Resource (ClinGen) ALS expert panel and classified as having evidence of disease association (Table 1) [8].

**Table 1:**
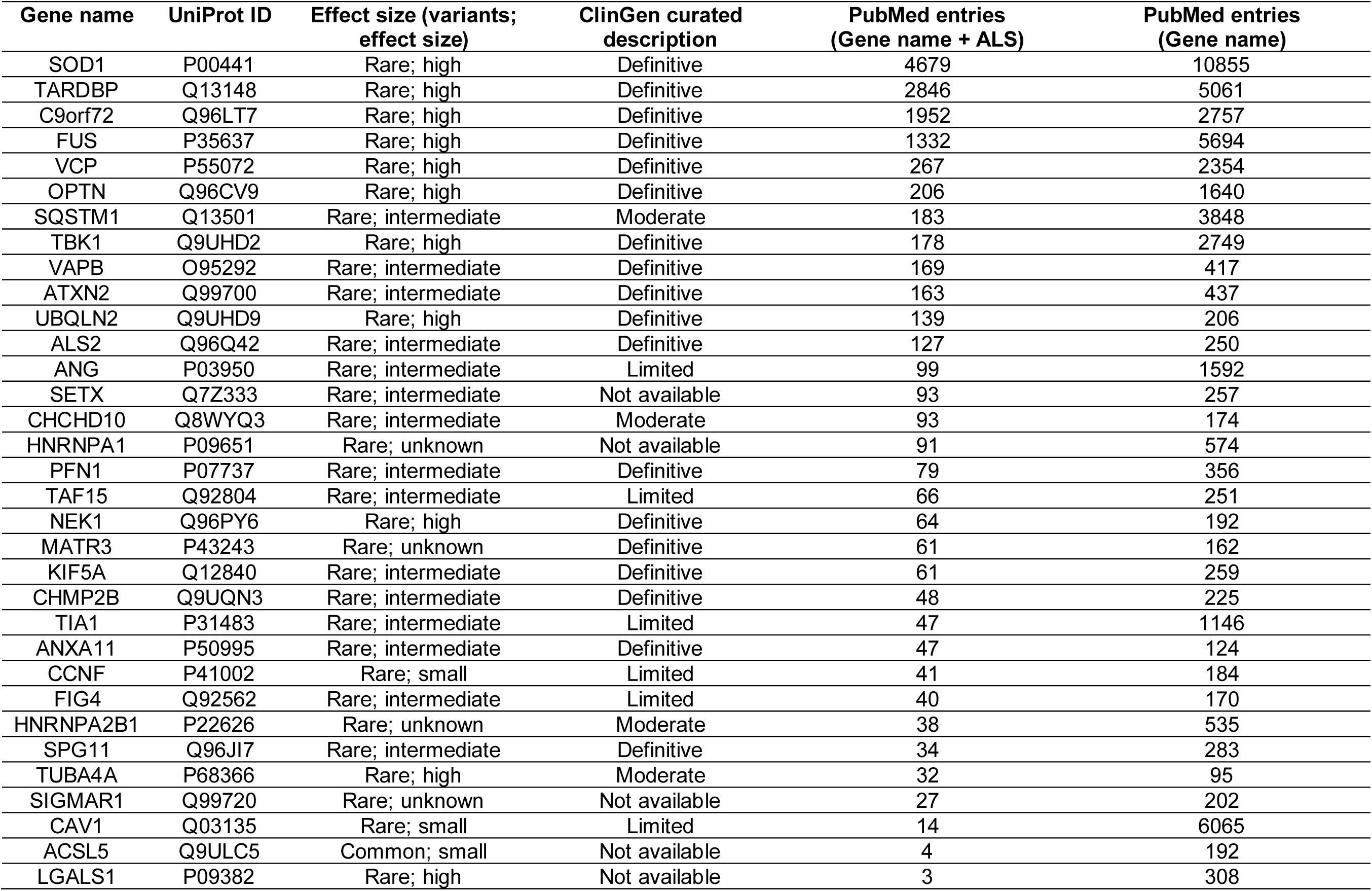

| Gene name | UniProt ID | Effect size (variants;<br>effect size) | ClinGen curated<br>description | PubMed entries<br>(Gene name + ALS) | PubMed entries<br>(Gene name) |
| --- | --- | --- | --- | --- | --- |
| SOD1 | P00441 | Rare; high | Definitive | 4679 | 10855 |
| TARDBP | Q13148 | Rare; high | Definitive | 2846 | 5061 |
| C9orf72 | Q96LT7 | Rare; high | Definitive | 1952 | 2757 |
| FUS | P35637 | Rare; high | Definitive | 1332 | 5694 |
| VCP | P55072 | Rare; high | Definitive | 267 | 2354 |
| OPTN | Q96CV9 | Rare; high | Definitive | 206 | 1640 |
| SQSTM1 | Q13501 | Rare; intermediate | Moderate | 183 | 3848 |
| TBK1 | Q9UHD2 | Rare; high | Definitive | 178 | 2749 |
| VAPB | O95292 | Rare; intermediate | Definitive | 169 | 417 |
| ATXN2 | Q99700 | Rare; intermediate | Definitive | 163 | 437 |
| UBQLN2 | Q9UHD9 | Rare; high | Definitive | 139 | 206 |
| ALS2 | Q96Q42 | Rare; intermediate | Definitive | 127 | 250 |
| ANG | P03950 | Rare; intermediate | Limited | 99 | 1592 |
| SETX | Q7Z333 | Rare; intermediate | Not available | 93 | 257 |
| CHCHD10 | Q8WYQ3 | Rare; intermediate | Moderate | 93 | 174 |
| HNRNPA1 | P09651 | Rare; unknown | Not available | 91 | 574 |
| PFN1 | P07737 | Rare; intermediate | Definitive | 79 | 356 |
| TAF15 | Q92804 | Rare; intermediate | Limited | 66 | 251 |
| NEK1 | Q96PY6 | Rare; high | Definitive | 64 | 192 |
| MATR3 | P43243 | Rare; unknown | Definitive | 61 | 162 |
| KIF5A | Q12840 | Rare; intermediate | Definitive | 61 | 259 |
| CHMP2B | Q9UQN3 | Rare; intermediate | Definitive | 48 | 225 |
| TIA1 | P31483 | Rare; intermediate | Limited | 47 | 1146 |
| ANXA11 | P50995 | Rare; intermediate | Definitive | 47 | 124 |
| CCNF | P41002 | Rare; small | Limited | 41 | 184 |
| FIG4 | Q92562 | Rare; intermediate | Limited | 40 | 170 |
| HNRNPA2B1 | P22626 | Rare; unknown | Moderate | 38 | 535 |
| SPG11 | Q96JI7 | Rare; intermediate | Definitive | 34 | 283 |
| TUBA4A | P68366 | Rare; high | Moderate | 32 | 95 |
| SIGMAR1 | Q99720 | Rare; unknown | Not available | 27 | 202 |
| CAV1 | Q03135 | Rare; small | Limited | 14 | 6065 |
| ACSL5 | Q9ULC5 | Common; small | Not available | 4 | 192 |
| LGALS1 | P09382 | Rare; high | Not available | 3 | 308 |

To estimate how ALS-associated proteins are represented in the literature, PubMed searches were performed for each gene using the gene name alone or combined with “ALS” (Table 1, ordered by number of PubMed entries). As shown in a line-scan plot (Figure S1), four genes—*SOD1*, *TARDBP*, *C9orf72*, and *FUS*—dominate the ALS literature, accounting for 10,809 of 13,173 publications (>80%). In contrast, the remaining 29 genes are far less studied in ALS and collectively define the “dark ALS-ome,” referring to ALS-associated proteins with limited functional characterization. Notably, seven of these genes—*VCP*, *OPTN*, *SQSTM1*, *TBK1*, *ANG*, *TIA1*, and *CAV1*—have been investigated in other biological contexts.

### Reagent evaluation and antibody development for ALS-RAP targets

According to the CiteAb database (citeab.com), only three of the 33 ALS-RAP targets have a top-cited renewable antibody characterized using genetic knockdown or KO approaches, highlighting a substantial gap in validated reagents. We therefore assessed renewable antibody availability across all targets prior to benchmarking with YCharOS characterization workflows, as batch-to-batch variability of polyclonal antibodies limits reproducibility [6, 14]. Six proteins—ANXA11, OPTN, MATR3, PFN1, UBQLN2 and VCP—had limited commercially renewable antibodies. To address this gap, the SGC generated recombinant single-chain variable fragment (scFv) antibodies optimized to recognize native, folded proteins, making them particularly well suited for IP [15]. Recombinant full-length proteins or protein domains were produced, purified, site-specifically biotinylated, and used as antigens for phage-display selections. Individual scFv clones were initially screened by ELISA, homogeneous time-resolved fluorescence, and suspension bead assays to confirm target binding and specificity. Candidate binders were subsequently evaluated by immunoprecipitation coupled to mass spectrometry (IP-MS) using established protocols [16], with at least three scFv clones screened for each target. The top-performing clone, selected based on IP-MS target enrichment, was then benchmarked alongside commercially available antibodies in the YCharOS validation pipeline. Antibody-encoding plasmids have been deposited at Addgene to ensure community access, and the amino acid sequences of reported scFvs are provided in Table S2.

In parallel, we evaluated KO cell line availability for antibody characterization. Twenty-one commercial KO lines were available from YCharOS partners and were selected based on parental expression levels above 2.5 log₂(TPM+1) in the DepMap database, a threshold suitable for antibody characterization studies [9]. To complete coverage, 12 additional custom KO lines were generated and are available upon request, while *VCP* was targeted with siRNA as it is an essential gene.

### Antibody performance and coverage for ALS-RAP proteins

Antibody performance for ALS-RAP targets was systematically evaluated in western blot (WB), IP, and immunofluorescence (IF) using KO-based characterization protocols [9]. In total, 303 antibodies (297 commercial and 6 custom scFv antibodies) targeting 33 ALS-associated proteins were tested. All antibodies were assessed in WB. For five proteins—ALS2, FIG4, NEK1, SIGMAR1, and SPG11—the absence of a specific antibody precluded confident assessment of antibody performance, as we could not determine whether this reflected uniformly poor antibody performance or suboptimal IF conditions for these targets. These datasets were therefore not included in Zenodo reports. Antibodies against ANG, CAV1, CCNF, KIF5A, and SETX were not evaluated in IP and/or IF due to the recent availability of renewable reagents, specialized experimental requirements (e.g., CCNF and cell-cycle arrest), or the secreted nature of ANG (Figure 1A).

**Figure 1:**
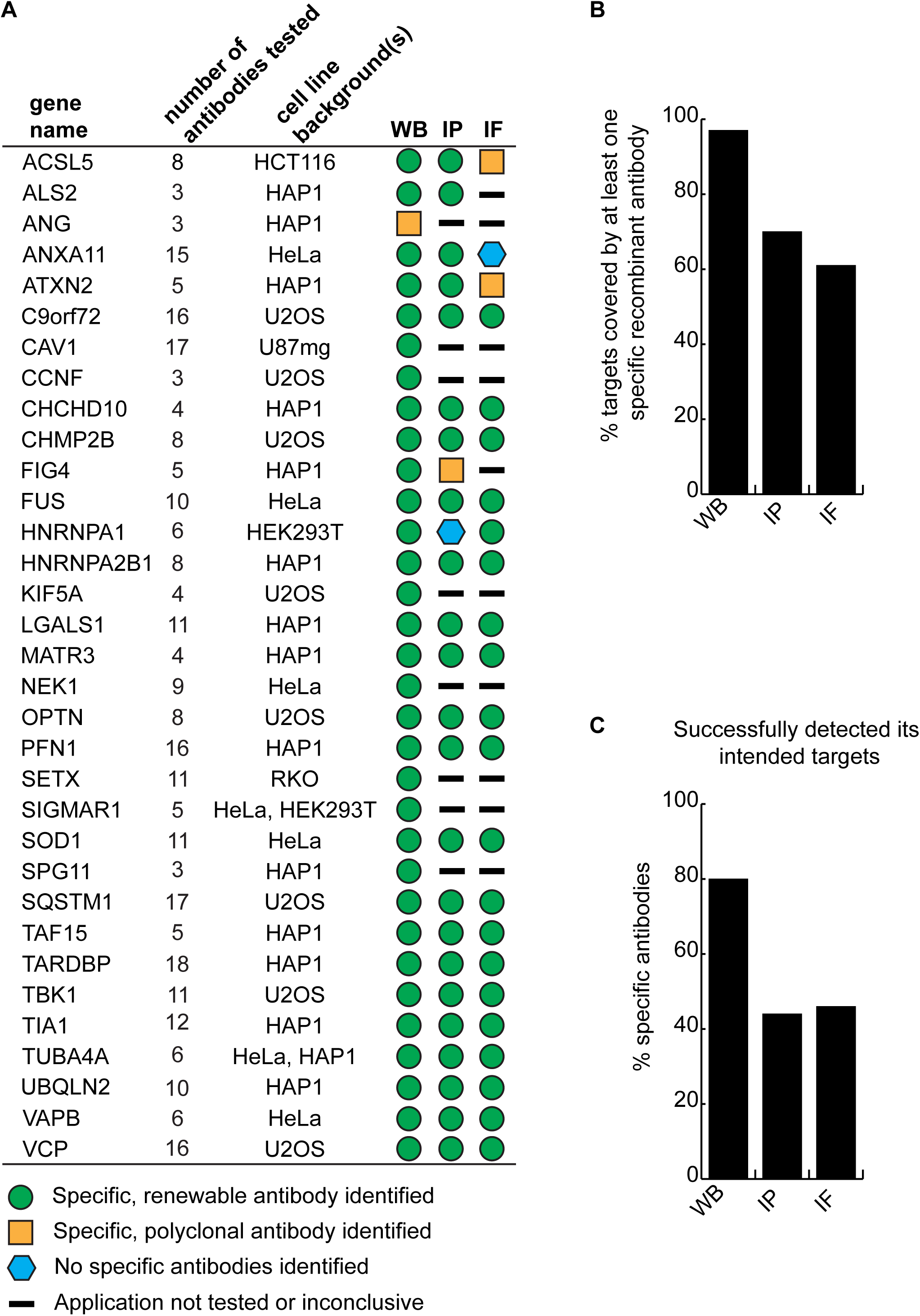
Summary of antibody performance for ALS-RAP proteins. **(A)** Overview of the identification of high-quality antibodies targeting ALS-RAP proteins. **(B)** Percentage of ALS-RAP targets covered by at least one specific renewable (monoclonal or recombinant) antibody for the indicated application. **(C)** Percentage of tested antibodies that met specificity criteria in each indicated application.

Antibodies were evaluated independently for each application as previously described [6]. In WB, antibodies were considered specific when an immunoreactive band was absent in the corresponding KO lysate. In IP, antibodies were considered successful when the target protein was specifically enriched relative to the starting material, as confirmed by immunoblot detection of the immunoprecipitated protein. In IF, antibodies were considered specific when the mean fluorescence intensity in wild-type cells was at least 1.5-fold higher than that measured in KO cells using the mosaic WT/KO assay. For IF, a minimum of 250 WT and 250 KO cells were quantified for each antibody. Detailed experimental workflows have been described previously [9].

Across all 33 targets, at least one renewable, KO-validated antibody was identified for 32 (97%), 23 (70%), and 20 (61%) proteins in WB, IP, and IF, respectively (Figure 1B). Overall, 243 of 303 antibodies (80%) detected their intended target in WB, 105 of 238 (44%) immunoprecipitated their target, and 107 of 231 (46%) demonstrated specific target staining in IF (Figure 1C).

The newly developed scFv antibodies filled key gaps; for example, the scFv against PFN1 was the only antibody—among 16 tested—capable of immunoprecipitating the protein [17]. Collectively, these results demonstrate that renewable, KO-validated antibodies cover most ALS-associated proteins, enabling reproducible studies of ALS biology.

For transparency and reuse, we generated 33 antibody characterization reports (one per protein), reviewed with YCharOS industry partners and deposited on Zenodo, with a subset published through the F1000Research YCharOS Gateway [18]. All antibodies tested are listed in Table S1 with links to their corresponding characterization reports. Research Resource Identifiers (RRIDs) were obtained for all antibodies and KO cell lines to facilitate reagent traceability [19, 20]. Antibodies used for downstream protein detection analyses were selected based on these characterization data, prioritizing renewable reagents. Catalogue numbers for these antibodies are provided in the Key Resources Table.

### Antibody characterization case study: Galectin-1

To illustrate the YCharOS antibody characterization workflow, we present the characterization of antibodies targeting Galectin-1, encoded by the *LGALS1* gene. Galectin-1 is a lectin implicated in immune and stress responses in the nervous system and has been reported in association with ALS in transcriptomic and functional studies [21, 22]. Antibodies were not assigned scores, consistent with the YCharOS philosophy that end users should evaluate raw data to select fit-for-purpose reagents based on their specific experimental needs.

Using DepMap transcriptomic data, the HAP1 cell line was selected for antibody characterization because it expresses *LGALS1* at adequate levels (7.9 log₂(TPM+1)), above the 2.5 threshold used for antibody characterization studies [6]. An *LGALS1* KO HAP1 cell line was obtained from Revvity, and ten renewable commercial antibodies from YCharOS partners were assembled for benchmarking (Table 2).

**Table 2:**

| Company | Catalog number | Lot number | RRID<br>(Antibody<br>Registry) | Clonality | Clone ID | Host | Concentration<br>(µg/µL) | Vendors<br>recommended<br>applications |
| --- | --- | --- | --- | --- | --- | --- | --- | --- |
| Abcam | ab108389** | 1018000-29 | AB_10863417 | recombinant<br>mono | EPR3205 | rabbit | 1.35 | WB, IF |
| Abcam | ab138513* | 1086382-8 | AB_2894851 | monoclonal | EPR3206(2) | rabbit | 0.26 | WB , IF |
| Abcam | ab192842** | 1011456-2 | AB_3725515 | recombinant<br>mono | SP247 | rabbit | 0.04 | WB |
| ABclonal | A21218** | 3522042922 | AB_3725523 | recombinant<br>mono | ARC53156 | rabbit | 1.20 | WB |
| Aviva Systems<br>Biology | ARP58491_P050 | QC25810-42978 | AB_2046045 | polyclonal | - | rabbit | 0.50 | WB |
| Bio-Techne<br>(R&D Systems) | MAB1152* | CJKC0325011 | AB_3657585 | monoclonal | 933133 | mouse | 0.50 | WB |
| Cell Signaling<br>Technology | 12936** | 3 | AB_2798065 | recombinant<br>mono | D608T | rabbit | 0.10 | WB , IF |
| Thermo Fisher<br>Scientific | MA5-32779** | 79527780 | AB_2810056 | recombinant<br>mono | JM13-37 | rabbit | 1.00 | WB , IP, IF |
| Thermo Fisher<br>Scientific | MA5-35566** | 79531623 | AB_2849466 | recombinant<br>mono | 7W5Q6 | rabbit | 2.00 | WB , IF |
| Thermo Fisher<br>Scientific | MA5-47904** | 79531680 | AB_3074348 | recombinant<br>mono | SAIC-25B-112 | rabbit | 1.00 | WB |

Antibody performance was evaluated across WB, IP, and IF using WT and KO HAP1 cells. In WB experiments, high-quality antibodies detected Galectin-1 in WT lysates with complete signal loss in KO lysates (Figure 2A). Galectin-1 was also detected in conditioned medium, consistent with its known secretion (Figure 2B). In IP experiments, antibodies were assessed for their ability to enrich Galectin-1 from cell lysates, with enrichment in the IP fraction and partial depletion from the unbound fraction indicating effective capture (Figure 2C). Finally, antibody performance in IF was evaluated using a mosaic strategy in which fluorescently labeled WT and KO cells were mixed and imaged in the same field of view, minimizing staining and imaging bias (Figure 2D). Complete experimental datasets and antibody characterization reports are available through Zenodo.

**Figure 2:**
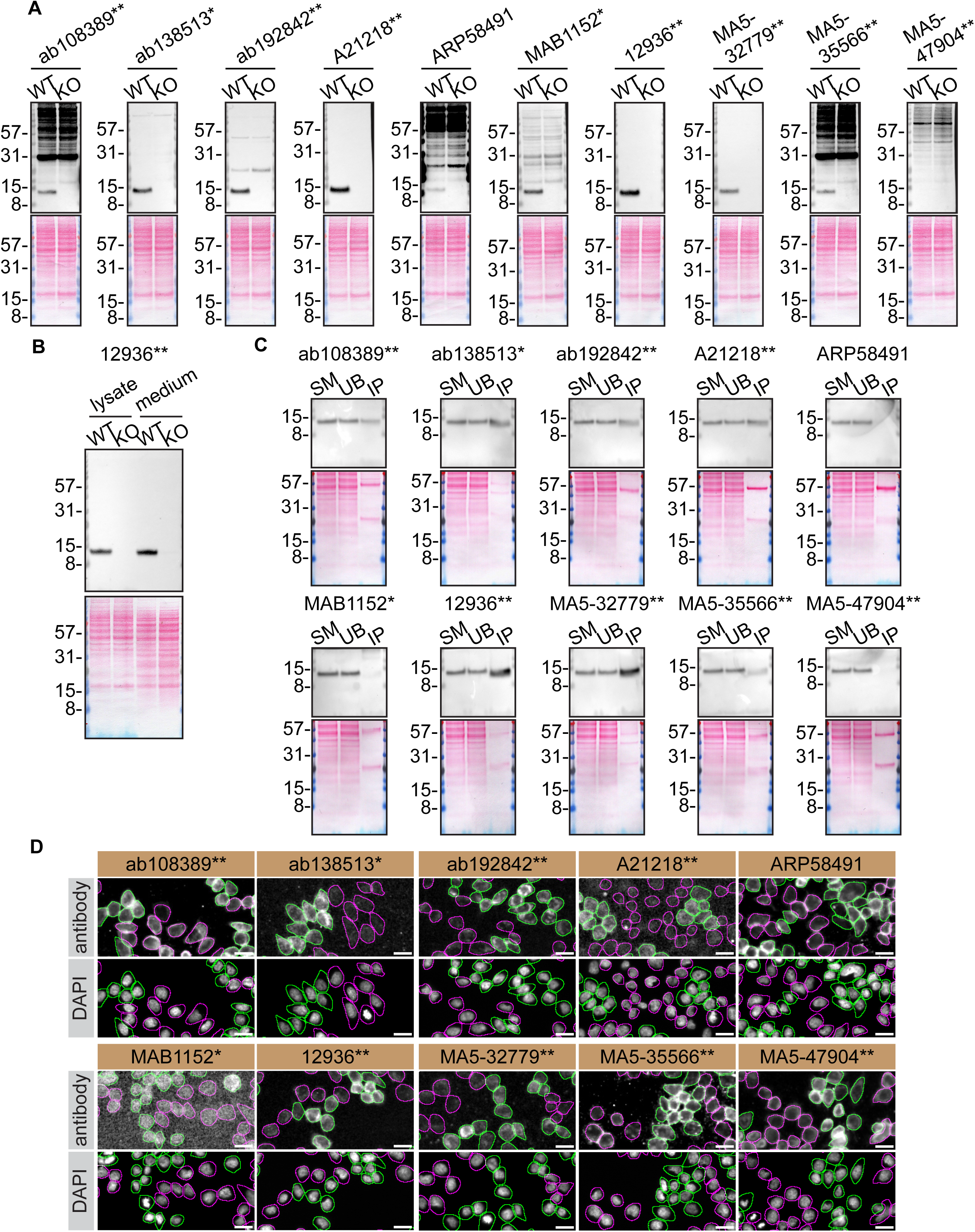
YCharOS antibody characterization workflow for Galectin-1 (*LGALS1*). Ten Galectin-1 antibodies were evaluated side-by-side across three applications using HAP1 WT and *LGALS1* KO cells. **(A)** Whole-cell protein lysates from HAP1 WT and *LGALS1* KO cells (20 µg per lane) were analyzed in WB using the indicated Galectin-1 antibodies. Ponceau-stained membranes are shown to assess protein loading and transfer efficiency. **(B)** Whole-cell protein lysates (10 µg) and conditioned media (10 µg) from HAP1 WT and *LGALS1* KO cells were analyzed in WB using antibody 12936, as in (A), to detect intracellular and secreted Galectin-1. **(C)** Immunoprecipitation was performed from HAP1 WT cell lysates using the indicated Galectin-1 antibodies. Starting material (SM), unbound fractions (UB), and whole immunoprecipitates (IP) were analyzed by WB. Anti-Galectin-1 antibody A21218 was used as a primary antibody. **(D)** HAP1 WT and *LGALS1* KO cells were differentially labeled with green and far-red fluorescent dyes, respectively, mixed at a 1:1 ratio, and plated in optically clear 96-well plates. Cells were stained with the indicated Galectin-1 antibodies, followed by Coralite 555–conjugated secondary antibodies and DAPI. Images were acquired in the blue (DAPI), green (WT), red (antibody staining), and far-red (KO) channels. Representative grayscale images of the blue and red channels are shown; WT and KO cells are outlined with green and magenta dashed lines, respectively. Scale bar, 10 µm. Recombinant antibodies are denoted by double asterisks (**) and monoclonal antibodies by single asterisks (*) after the catalogue numbers; polyclonal antibodies are unmarked. Antibody screening was performed in two independent experiments; representative data from one experiment are shown.

### ALS-RAP antibody recommendations on the OGA website

In addition to dissemination through YCharOS, ALS-RAP antibody characterization datasets were independently evaluated by the Only Good Antibodies (OGA) initiative [2], which did not participate in generating the experimental data. OGA integrates KO-based characterization datasets from YCharOS into a publicly accessible antibody assessment platform (https://onlygoodantibodies.co.uk/), where reagents are curated and presented to facilitate antibody selection by end users. Antibody entries include direct links to manufacturer purchasing pages. A screenshot of the Galectin-1 dataset illustrates the interface: a schematic representing high-quality antibodies is shown at the top, while all tested antibodies are listed below with application-specific recommendations derived from OGA analysis of the YCharOS data (Figure 3).

**Figure 3:**
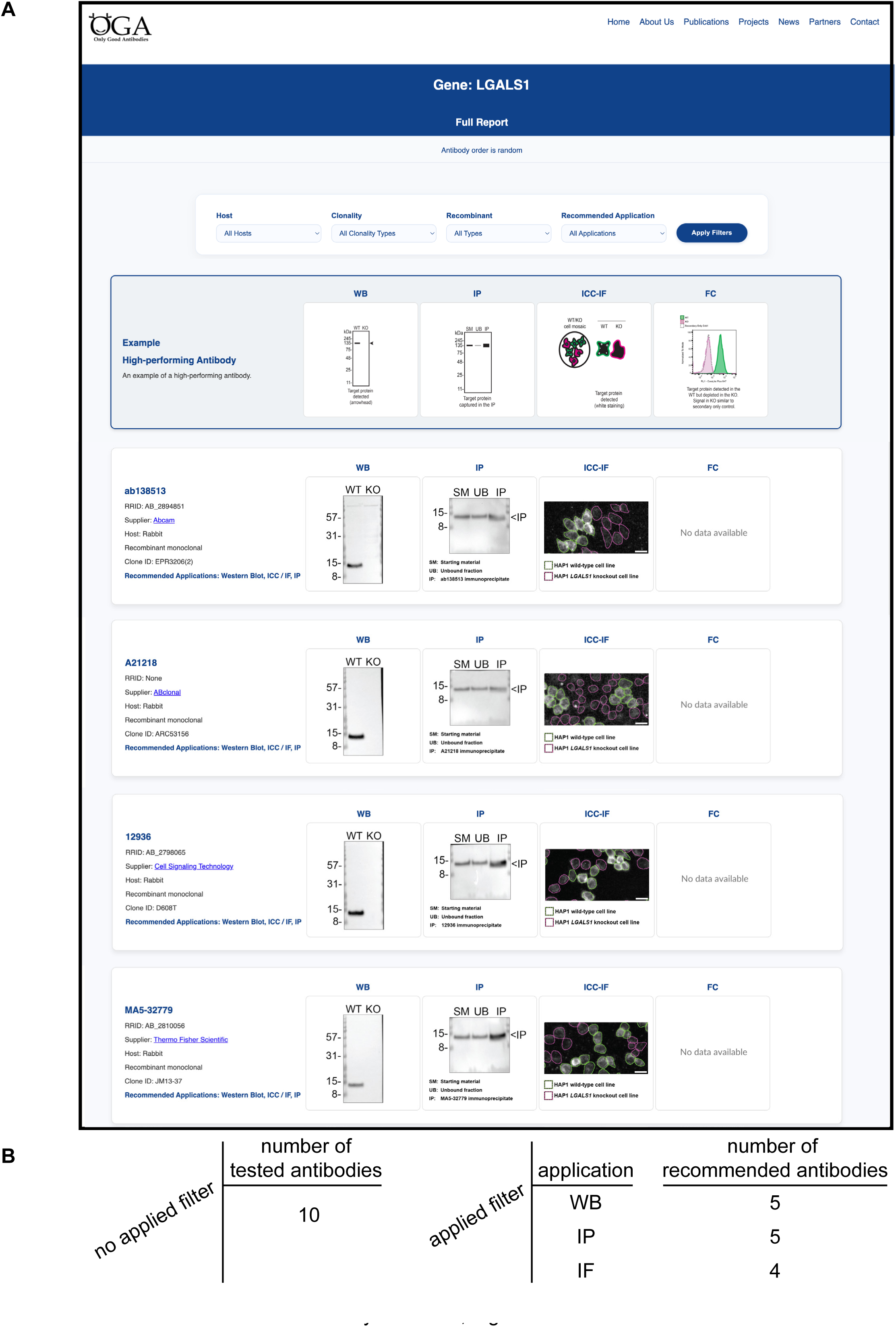
Accessing curated antibody characterization data through the OGA website. YCharOS antibody characterization data for ALS-RAP proteins are accessible through the Only Good Antibodies (OGA) website (https://onlygoodantibodies.co.uk/). The OGA homepage provides a searchable interface for the 33 ALS-RAP proteins, together with 35 additional neurological disease–associated proteins evaluated by YCharOS, and includes links to key OGA publications, data interpretation guidelines, antibody characterization protocols, and a portal for nominating new targets. **(A)** Representative results page generated by searching *LGALS1*, showing filters for host species, clonality, recombinant format, and OGA-recommended applications; the WB filter is applied. A schematic panel at the top illustrates ideal antibody performance across applications, while antibodies matching the selected filters are displayed in randomized order. **(B)** Number of antibodies returned following application of the selected filters.

By adding an interpretive layer to the raw characterization datasets, OGA improves the discoverability of validated antibodies against ALS-associated proteins while preserving full access to the underlying data. This integration complements the ALS-RAP resource by pairing systematic KO-based characterization with a user-oriented recommendation framework.

### Neurological expression landscape of ALS-RAP proteins

#### Cellular context for ALS-RAP expression analysis

To provide context for protein expression analyses, we first examined RNA expression of ALS-associated genes using the Human Protein Atlas (HPA) single-nucleus brain dataset [23]. Genes were categorized as neuron-enriched, glial-enriched (astrocytes, oligodendrocytes, and microglia), or broadly expressed based on relative RNA enrichment across cell types. Several genes showed preferential neuronal expression (*ALS2*, *CCNF*, *KIF5A*, and *TUBA4A*), whereas others were enriched in glial populations (*ACSL5*, *ANXA11*, *C9orf72*, *CAV1*, *CHMP2B*, *FIG4*, *FUS*, *HNRNPA1*, *LGALS1*, *SETX*, *SQSTM1*, and *TBK1*). The remaining genes were broadly expressed, while four (*ANG*, *MATR3*, *SIGMAR1*, and *TIA1*) were barely detected across neurological cell types (Figure S2). Together with previously published single-cell RNA sequencing datasets of human brain [24], these analyses highlight the heterogeneous expression of ALS-associated genes.

Because RNA abundance does not necessarily predict protein levels [25, 26], we next examined ALS-RAP protein expression across neurological cell types. Human iPSC-derived motor neurons, dopaminergic neurons, oligodendrocytes, astrocytes, and microglia were analyzed, complemented with primary human fetal astrocytes, fetal microglia, and monocyte-derived macrophages (MDMs). Cell identity was confirmed using established marker analyses (Figures S3–S4).

#### Protein profiling across neurological cell types

We profiled 30 ALS-RAP proteins in WB using two independent preparations of the same cell types: one analyzed by chemiluminescence-based detection and the other by fluorescence-based WB (Figure 4A-C). For each target, the best-performing renewable antibody identified during YCharOS screening was used. KO-validated primary bands were consistently detected at the expected molecular weight across neurological cell types. Additional bands were occasionally observed (e.g., MATR3, SQSTM1, ANXA11, ACSL5, UBQLN2) but were not interpreted, as we could not determine whether they corresponded to the intended target or reflected antibody cross-reactivity. Three proteins were excluded from expression profiling: CCNF (requires cell-cycle arrest for detection), ANG (secreted protein not readily detected in lysates), and SETX (insufficient signal-to-noise). Relative signal intensities within each detection modality were compared to identify general patterns of protein detection across neuronal and glial populations.

**Figure 4:**
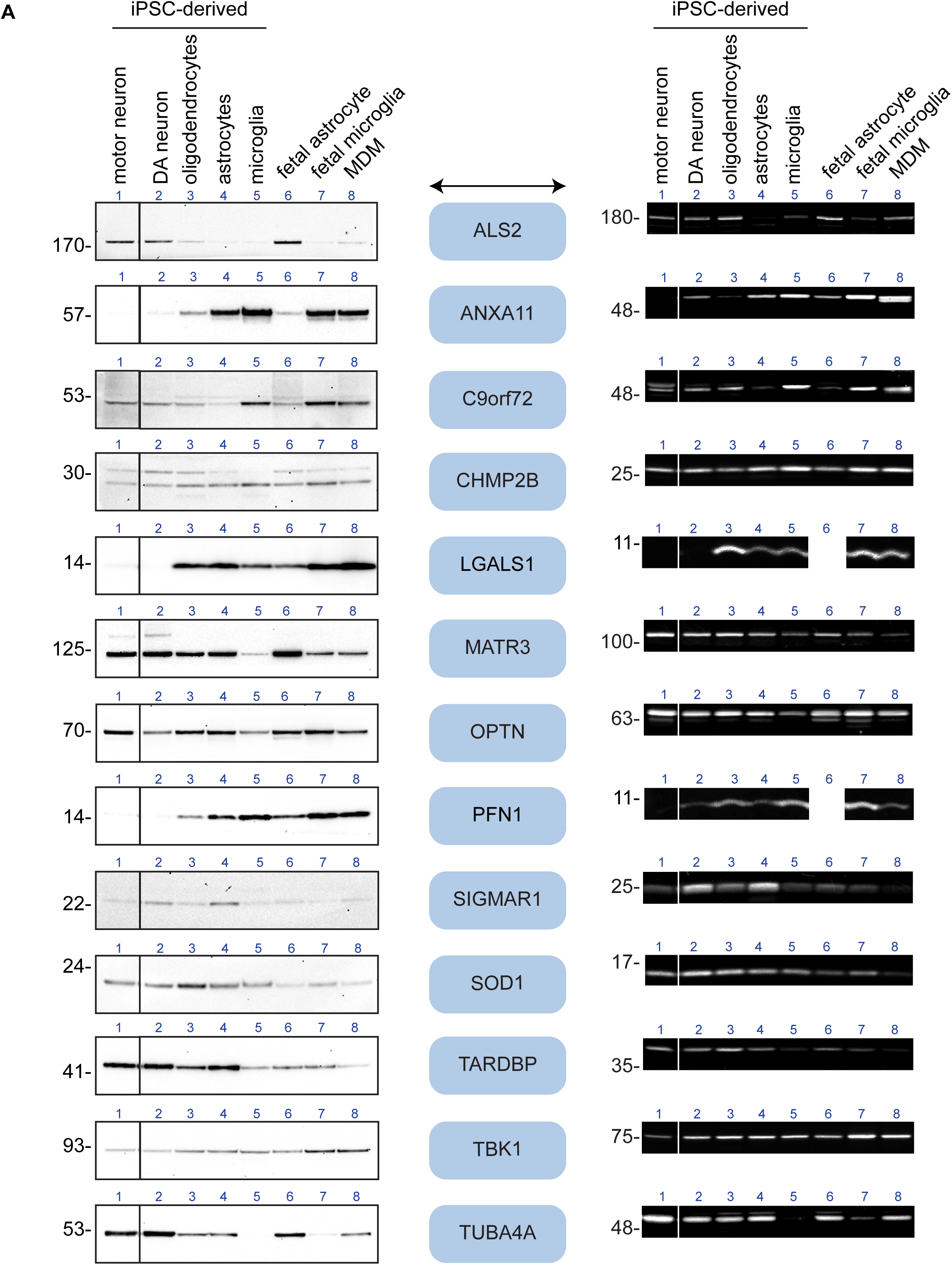
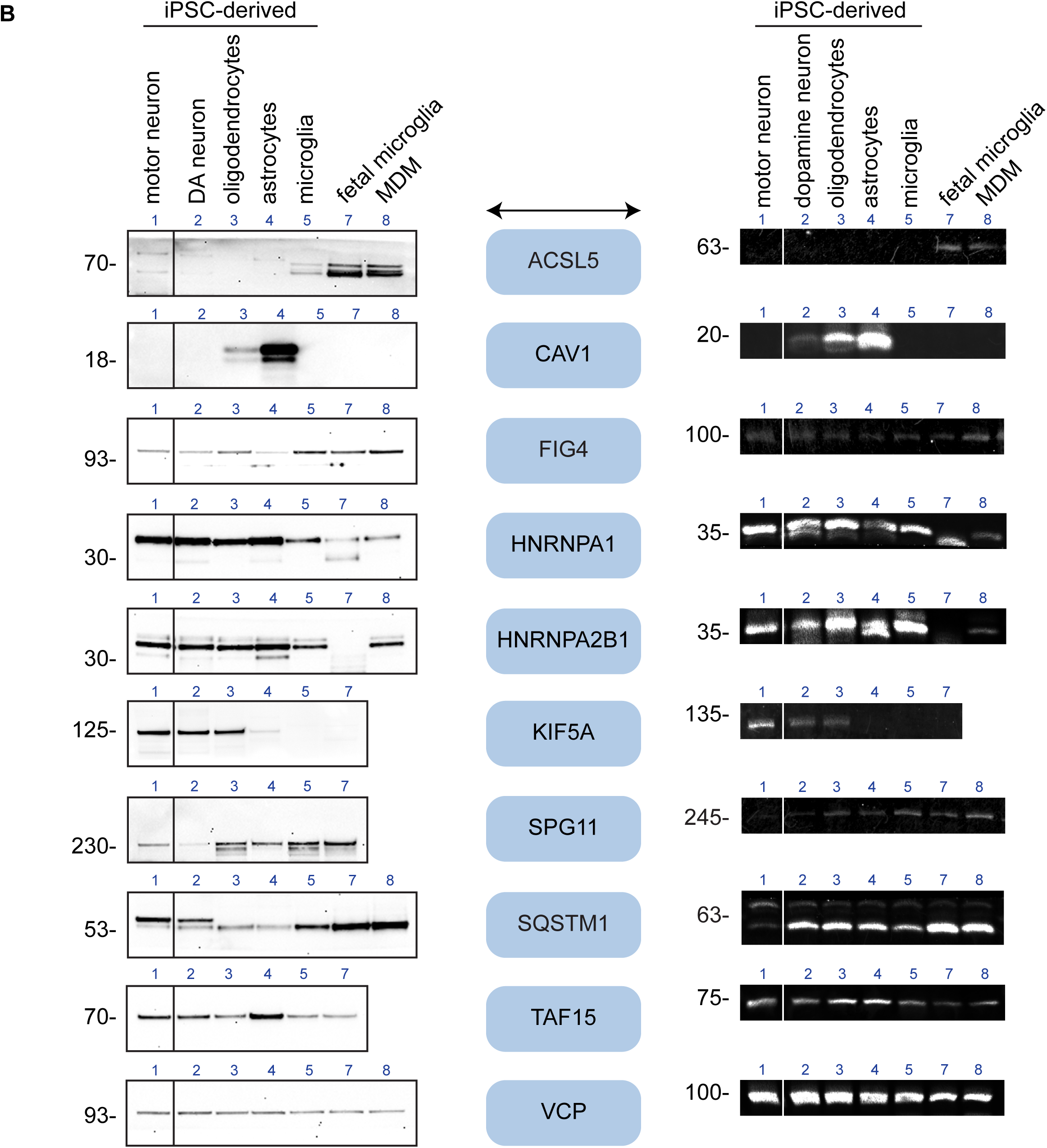
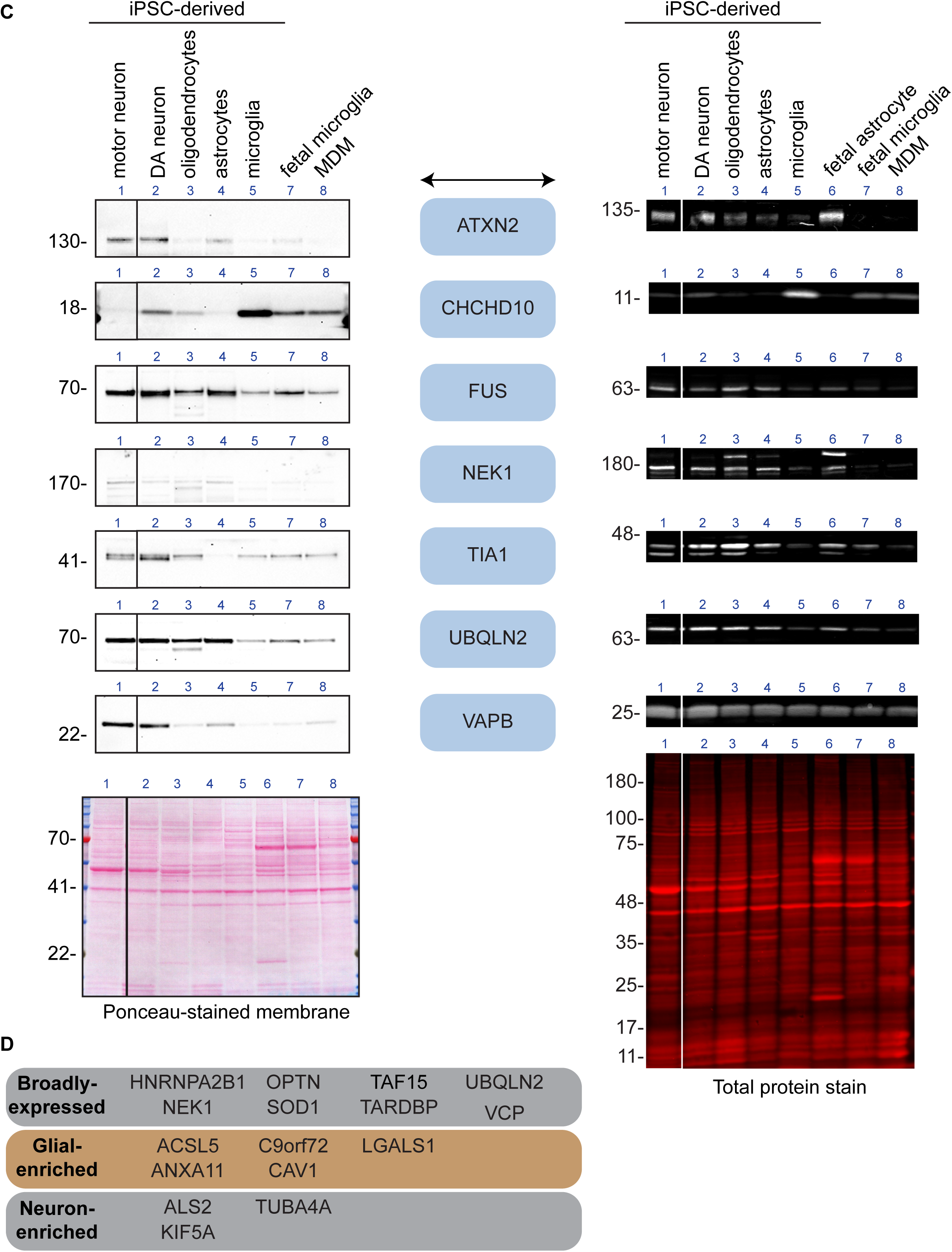
Detection of 30 ALS-RAP proteins across human-derived neurological cell types. WB showing detection of 30 ALS-RAP proteins across the indicated cell types. For each protein, two independent experiments are shown: chemiluminescence-based WB (left), performed using a defined lysate batch across all corresponding blots, and fluorescence-based WB (right), performed using a second independent lysate batch. Lysates were prepared from the indicated human iPSC-derived cell types and primary cells. Proteins are listed in alphabetical order. A representative Ponceau-stained membrane and total protein stain are shown to assess protein loading and transfer efficiency. Vertical lines between the motor neuron and DA neuron lanes indicate that the blots were spliced to remove additional conditions not relevant to this study. In **(A)**, eight lysates were used across both detection modalities. In **(B)**, the fetal astrocyte lysate was not available for either detection modality. In **(C)**, the fetal astrocyte lysate was not included in the chemiluminescence-based experiment. Lane identities: 1, motor neurons; 2, DA neurons; 3, oligodendrocytes; 4, astrocytes; 5, microglia, 6, fetal astrocytes; 7, fetal microglia; 8, MDMs. DA, dopaminergic; MDM, monocyte-derived macrophages. **(D)** Summary of 16 targets for which WB data and Human Protein Atlas RNA expression profiles are concordant, grouped by expression category.

Many ALS-RAP proteins were detected across both neuronal and glial cell types, including ALS2, CHMP2B, FIG4, FUS, HNRNPA1, HNRNPA2B1, MATR3, NEK1, OPTN, SIGMAR1, SOD1, SQSTM1, TAF15, TARDBP, TBK1, TIA1, UBQLN2, and VCP. Several proteins showed higher signal in glial populations in both datasets, including ACSL5, ANXA11, C9orf72, CAV1, CHCHD10, LGALS1, PFN1, and SPG11. ACSL5, ANXA11, C9orf72, and CHCHD10 were detected at higher levels in microglial populations, while CAV1 showed stronger signal in astrocytes. These patterns were consistent between iPSC-derived and primary glial cells. In contrast, several proteins showed higher protein levels in neuronal populations. ATXN2 was strongly detected in motor and dopaminergic neurons, while KIF5A and TUBA4A were enriched in neuronal lineages. VAPB was primarily detected in neurons, with additional signal in glial cells in the fluorescence-based dataset. Overall, these analyses reveal diversity in the cellular distribution of ALS-RAP proteins. Notably, many ALS-associated proteins were detected across glial and immune cell populations, supporting the view that ALS genetics converges on multicellular disease mechanisms.

Overall, 16 of 27 proteins assessed at both the RNA and protein levels showed concordant cell type categorization (Figure 4D), highlighting both concordance and divergence between transcriptomic and protein-level datasets. Consistent with the limited protein-level characterization of ALS-associated genes, STRING network analysis revealed sparse evidence of direct interactions among the 33 proteins, with clusters primarily involving autophagy regulators (TBK1, OPTN, SQSTM1) and RNA-binding proteins (FUS, TARDBP, MATR3, HNRNPA1, HNRNPA2B1) (Figure S5). These clusters mainly comprise broadly expressed proteins, whereas interactions involving cell type-restricted ALS-RAP proteins remain largely unexplored.

## Discussion

Large-scale genetic studies have identified hundreds of risk loci across neurodegenerative diseases, including Alzheimer’s disease [27, 28], Parkinson’s disease [29, 30], and ALS [31]. Despite these advances, functional follow-up has lagged behind genetic discovery, in part because disease-associated proteins lack validated reagents and systematic protein-level characterization. As a result, mechanistic insight has remained concentrated on a limited subset of genes, while the majority of risk-associated proteins remain poorly studied [32, 33]. In ALS, this gap is particularly evident, with many proteins remaining part of the “dark ALS-ome”.

Here, we address this gap through ALS-RAP, which provides a publicly available dataset of KO-characterized antibodies across three commonly used applications—WB, IP, and IF—enabling the selection of fit-for-purpose reagents for studying ALS-associated proteins. Using these reagents, we examined protein levels across iPSC-derived and primary neurological cell types, revealing diverse expression patterns, including frequent detection of ALS-associated proteins in glial populations, particularly microglia and macrophages.

Consistent with this gap, STRING network analysis revealed sparse connectivity among ALS-associated proteins. Our expression analyses identify cell type-specific patterns that may help contextualize potential biological relationships. For example, ANXA11, classified as a definitive ALS gene by ClinGen, was predominantly detected in microglial populations. Annexins regulate membrane dynamics and trafficking, yet the role of Annexin A11 in microglia or macrophages remains largely unexplored. Its enrichment in microglia places it in the same cellular context as several ALS-associated proteins involved in autophagy and vesicular pathways, including C9orf72, TBK1, UBQLN2, SQSTM1, and OPTN, supporting potential convergence on shared pathways in myeloid-lineage cells. C9orf72 also showed higher expression in microglial populations, in agreement with studies implicating C9orf72 in microglial functions such as phagocytosis and immune regulation [34, 35], as well as evidence that microglia present C9orf72-derived peptides capable of activating CD4⁺ T cells [36]. Together, these findings support the view that ALS genetics converges on multicellular disease mechanisms involving both neuronal and glial populations.

## Materials and Methods

### Analysis of PubMed entries for each ALS-associated gene

Throughout the manuscript, we reference ALS-RAP targets using their official gene names as listed in UniProt, written in italics (e.g., *SOD1*). To refer to the corresponding proteins, we use the same gene names in regular font, as the official protein names are often too long or ambiguous for practical use.

Each target gene was queried in PubMed using two search strategies: (1) GENE_NAME[Title/Abstract] to retrieve the total number of publications associated with the gene, and (2) (GENE_NAME[Title/Abstract]) AND (“amyotrophic lateral sclerosis” [Title/Abstract]) to identify publications specifically linking the gene to ALS. Searches were performed on August 27, 2025.

### Evaluation of significance of ALS-associated genes

Each candidate gene was assessed for its potential association with ALS using the evidence published at the time of this manuscript. Using the recent review of Nijs & van Damme as a model [7], we used currently available evidence to prioritize genes for potential association with ALS in a similar manner. First, genes were qualitatively assessed for rarity of genetic variants in the general population and among individuals with ALS. Next, any reported odds ratios (OR) for risk of ALS in variant carriers were used to approximate whether genes were considered to have “small” (OR < 1.2), “intermediate” (1.2 < OR < 10), or “high” (OR > 10) effect sizes. Genes were assigned an “uncertain” OR estimate if no OR had been published at the time of analysis. Variant frequency was assessed considering the type(s) of inheritance previously described; variants reported through GWAS or genotyping studies were designated as “common” whereas variants reported in autosomal family/pedigree-based studies were designated as “rare”. If the method of discovery was poorly described or through non-genetic analyses (e.g. transcriptomic or proteomic), genes were described as having “uncertain” frequency.

### RNA expression analysis

We accessed the single-cell transcriptomics resource of the Human Protein Atlas (HPA), specifically the single-nuclei brain dataset which includes single-cell RNA sequencing data generated from 2.5 million cells and mapped to 34 supercluster cell types. Data were generated from post-mortem human brain samples digested to a single cell suspension, and cell types were clustered into transcriptomic cell types.[37] Raw sequencing data was normalized using Trimmed mean of M values and expression values were presented as normalized transcripts per million (nTPM). RNA expression data for all 33 ALS genes were downloaded online from the HPA. Data were grouped together based on overall cell type (neuron, oligodendrocyte, astrocyte, microglia, etc.) and the average expression in each cell type was determined for each gene. The average neuron, oligodendrocyte, astrocyte, and microglia expression data were extracted from the overall dataset and used in downstream analyses.

In our analyses, genes were considered “expressed” if their expression values in at least three cell types were greater than the median expression value of the entire HPA dataset (9.2 nTPM). Genes were considered “barely detectable” if expression was less than the median value in at least two cell types. For a small subset of “barely detectable” genes, expression in one cell type was greater than the median value and substantially “enriched” in that cell type. We included these in the category of “expressed” genes despite the low expression in most cell types.

To describe the cell type expression patterns for each gene, we established criteria determining if mRNA expression levels for a gene is “enriched” or not in a given cell type. Cell types in which mRNA expression for a gene is “enriched” were defined as when the ratio of the expression value in that cell type to the average expression value across all cell types was greater than or equal to 1.5*. Each gene was categorized based on the “enriched” cell types. If either more than one cell type was “enriched” or no cell types were, the gene was categorized as having “low cell-type specificity”.

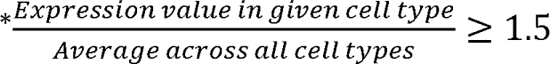

### String analysis

STRING analysis was performed by entering the UniProt IDs of the 33 ALS-RAP proteins into the “multiple proteins” search at string-db.org. The analysis was restricted to the “physical subnetwork”, with the meaning of network edges set at “confidence”, and the minimum required interaction score set to “highest confidence”.

### Western blot (WB)

Primary cells, iPSC-derived cultures and cultured cells were collected in RIPA buffer (25mM Tris-HCl pH 7.6, 150mM NaCl, 1 % NP-40, 1% sodium deoxycholate, 0.1% SDS) supplemented with protease inhibitors. Lysates were sonicated briefly for 5-10 secs depending on the volume. Following 30 min on ice, lysates were spun at ∼110,000 x *g* for 15 min at 4°C and equal protein aliquots of the supernatants were analyzed by SDS-PAGE and WB.

### Chemiluminescent WB

WB were performed with precast 10% Bis-Tris polyacrylamide gels (MOPS or MES SDS buffer) and nitrocellulose membranes. Total proteins were visualized with a Ponceau staining. Blots were blocked with 5% milk in TBS with 0.1% Tween 20 (TBST). Antibodies were also diluted in 5% milk in TBST and incubated O/N at 4°C. Membranes were washed three washes in TBST the next day. The peroxidase conjugated anti rabbit and anti-mouse secondary antibodies (Thermo Fisher Scientific) were diluted in 5% milk in TBST at 0.5 and 1 µg/mL, respectively. Secondaries were incubated with the membranes for 1 hr at room temperature followed by three washes. Membranes were incubated with ECL for 1 min prior to detection with the iBright CL1500 Imaging System (Thermo Fisher Scientific).

### Fluorescent WB

WB were performed with large 5–16% gradient Tris-based polyacrylamide gels (Tris-Glycine/SDS Buffer) and nitrocellulose membranes. Total proteins were visualized with the REVERT total protein stain (LI-COR Biosciences). Blots were blocked with 5% milk in TBST, and antibodies were diluted in the same buffer and incubated O/N at 4°C followed by three washes. The secondary antibodies (Odyssey IRDye 800CW or IRDye 680RD) were diluted in 5% milk in TBST to a final concentration of 0.05 µg/mL for anti-rabbit and 0.1 µg/mL for anti-mouse. Secondaries were incubated with the membranes for 1 hr at room temperature followed by three washes while covered from the light. Immuno-reactive bands were detected using the LI-COR Odyssey Imaging System (LI-COR Biosciences), and data analysis was done using LI-COR Image Studio Lite Version 6.10.2.

### Galectin-1 antibody screening in WB

For cell lysis, HAP1 WT and *LGALS1* KO cells were collected in RIPA buffer and processed as described above. To collect the conditioned medium, HAP1 WT and *LGALS1* KO cells were washed 3x with PBS 1x and starved for ∼36 hrs in Iscove’s Modified Dulbecco’s Medium (IMDM) supplemented with 1% Pen/Strep and 1% L-Glutamine. Culture media were collected and centrifuged for 10 min at 500 x *g* to eliminate cells and larger contaminants, then for 10 min at 4,500 x *g* to eliminate smaller contaminants. Culture media were then concentrated by centrifuging at 4,000 x *g* for 30 min using the Amicon Ultra-15 Centrifugal Filter Units with a membrane NMWL of 10kDa. Culture media were finally supplemented with 1x protease inhibitor cocktail mix.

10% Bis-Tris gels with MES buffer were used for SDS-PAGE and WB was performed using the primary antibodies (Table 2) at the following dilutions: ab108389 at 1/100, ab138513 atv1/10 000, ab192842 at 1/100, A21218 at 1/11 000, ARP58491 at 1/100, MAB1152 at 1/250, 12936 at 1/1000, MA5-32779 at 1/2000, MA5-35566 at 1/200, MA5-47904 at 1/1000. Anti-rabbit and anti-mouse antibodies (Proteintech) were used at a final concentration of 0.05 µg/ml and 0.5 µg/ml, respectively.

### Galectin-1 antibody screening in immunoprecipitation (IP)

Antibody-bead conjugates were prepared by adding 2 µg to 500 µl of Pierce IP Lysis Buffer from Thermo Fisher Scientific in a microcentrifuge tube, together with 30 µl of Dynabeads protein A-(for rabbit antibodies). Tubes were rocked for 1 h at 4°C followed by two washes to remove unbound antibodies. HAP1 WT were collected in Pierce IP buffer (25 mM Tris-HCl pH 7.4, 150 mM NaCl, 1 mM EDTA, 1% NP-40 and 5% glycerol) supplemented with protease inhibitor. Lysates were rocked 30 min at 4°C and spun at 110,000 x *g* for 15 min at 4°C. 0.2 ml aliquots at 1 mg/ml of lysate were incubated with each antibody-bead conjugate for 2 h at 4°C. To run 4% SM (8 µl per SM) alongside each IP on the gel, a master mix of lysate was kept aside on ice for subsequent SM sample preparation. The UBs were collected separately and prepared for gel loading similar to the rest of the samples. Beads were washed three times with 1.0 ml of IP buffer and processed for SDS-PAGE on precast midi 10% Bis-Tris polyacrylamide gels. For WB, membranes were trimmed at 24 kDa and the lower part was incubated with the rabbit recombinant monoclonal anti-Galectin-1 antibody A21218 diluted at 1/3000 in 5% milk in TBST. HRP-conjugated anti-rabbit (Proteintech) diluted in 5% milk in TBST was used as a secondary antibody at 0.05 µg/ml.

### Galectin-1 antibody screening in immunofluorescence (IF)

WT and *LGALS1* KO HAP1 cells were labelled with a green and a far-red fluorescence dye, respectively. WT and KO cells were plated 96-well plate with optically clear flat-bottom (Perkin Elmer) as a mosaic and incubated for 24 hrs in a cell culture incubator at 37°C, 5% CO2. Cells were fixed in 4% paraformaldehyde (PFA) in phosphate buffered saline (PBS). Cells were permeabilized in PBS with 0.1% Triton X-100 for 10 min at room temperature and blocked with PBS with 5% BSA, 5% goat serum and 0.01% Triton X-100 for 30 min at room temperature. Cells were incubated with IF buffer (PBS, 5% BSA, 0.01% Triton X-100) containing the primary Galectin-1 antibodies overnight at 4°C. Cells were then washed 3 × 10 min with IF buffer and incubated with the corresponding CoraLite Plus 555 secondary antibody in IF buffer at a dilution of 0.5 µg/ml for 1 hr at room temperature. Cells were washed 3 × 10 min with 1x PBS. The nuclei were labelled with DAPI fluorescent stain at a concentration of 5 ng/ml in 1x PBS for 3 min then washed once with 1x PBS.

Images were acquired on an ImageXpress micro confocal high-content microscopy system (Molecular Devices). The objective used is a water Apo LambdaS LWD with magnification of 20x, NA 0.95 and scientific CMOS cameras, equipped with 395, 475, 555 and 635 nm solid state LED lights (lumencor Aura III light engine) and bandpass filters to excite DAPI, Cellmask Green, Coralite-555 and Cellmask Red, respectively. Images had pixel sizes of 0.68 x 0.68 microns, and a z-interval of 4 microns. For analysis and visualization, shading correction (shade only) was carried out for all images. Then, maximum intensity projections were generated using 3 z-slices. Segmentation was carried out separately on maximum intensity projections of Cellmask channels using CellPose 1.0, and masks were used to generate outlines and for intensity quantification.

### iPSC differentiation protocols

Healthy control iPSCs from the AIW002-02 & QPN929 backgrounds (Early Drug Discovery Unit-EDDU, McGill University) were selected for differentiation into the indicated neurological cell types. In all cases, iPSCs were cultured in mTeSR1 for 5-7 days prior to differentiation. Two batches of neurological cells were generated from the AIW002-02 background using identical protocols, with the exception that the QPN929 iPSC line was used for microglia differentiation in batch used for chemiluminescent WB.

#### Motor neuron differentiation

Motor neurons were differentiated from iPSCs following a previously published protocol [38, 39]. Basic neural medium (BNM, 1:1 mixture of DMEM/F12 and Neurobasal, 0.5X N2, 0.5X B27, 0.5X GlutaMAX, 1X antibiotic-antimycotic, and 100 μM ascorbic acid) serves as the base medium for all other formulations used in this protocol. iPSCs were seeded on matrigel coated flasks in neural induction medium (BNM supplemented with 2 μM SB431542, 2 μM DMH1, 3 μM CHIR99021) with a full media change performed every other day. After 6 days, cells were passaged in patterning medium (BNM supplemented with 1 μM CHIR99021, 2 μM SB431542, 2 μM DMH1, 0.1 μM retinoic acid (RA) and 0.5 μM purmorphamine (PMN)) onto poly-L-ornithine- and laminin-coated flasks and cultured for a further 6 days with a full media change performed every other day. Cells were then passaged onto poly-L-ornithine- and laminin-coated flasks in expansion medium (BNM supplemented with 3 μM CHIR99021, 2 μM SB431542, 2 μM DMH1, 0.1 μM RA and 0.5 μM PMN, and 0.5 mM valproic acid (VPA), with a full media change performed every other day for 6 days to obtain motor neural progenitor cells (M-NPCs). M-NPCs were dissociated as single cells and seeded onto poly-L-ornithine- and laminin-coated plates in final differentiation medium (BNM supplemented with 0.5 μM RA, 0.1 μM PMN, 0.1 μM Compound E (CpdE), 10 ng/mL insulin-like growth factor-1 (IGF-1), brain-derived neurotrophic factor (BDNF), and ciliary neurotrophic factor (CNTF)), with a half media change once per week for final differentiation into motor neurons (MNs). MNs were considered mature and collected for WB after four weeks of differentiation.

#### Dopaminergic neuron differentiation

Dopaminergic neurons were differentiated from iPSCs following a protocol of floor-plate induction with additional modifications [40, 41]. Briefly, iPSCs were dissociated into single cells to form embryoid bodies (EBs). EBs were grown on low-attachment plates for one week in DMEM/F12 supplemented with 1X N2 and 1X B27, in the presence of 10 μM SB431542, 200 ng/mL Noggin, 200 ng/mL sonic hedgehog, 3 μM CHIR99021 and 100 ng/mL Fibroblast growth factor 8 (FGF8). On day 7, EBs were transferred, in the same media, to poly-L-ornithine- and laminin-coated plates. On day 14, rosettes were selected semi– manually and cultured as a monolayer in DMEM/F12 supplemented with 1X N2 and 1X B27 on poly-L-ornithine- and laminin-coated plates to generate dopaminergic neural progenitor cells (DA-NPCs). DA-NPCs were passaged at a ratio of 1:3 every 5-7 days. DA-NPCs were then cultured in neurobasal medium, supplemented with 1X N2 and 1X B27, in the presence of 1 μg/mL laminin, 500 μM dibutyryl-cAMP (db-cAMP), 20 ng/mL BDNF, 20 ng/mL glial-derived neurotrophic factor (GDNF), 200 μM ascorbic acid, 100 nM CpdE and 1 ng/mL transforming growth factor β3 (TGF-β3), with a half media change twice per week for final differentiation into dopaminergic neurons. Dopaminergic neurons were considered mature and collected for WB after four weeks of differentiation.

#### Oligodendrocyte differentiation

Oligodendrocytes were differentiated from iPSCs following a recently published protocol [42]. The base media for all other formulations in this protocol is as follows: DMEM/F12 with Glutamax supplemented with 1X N2, 1X B27, 1X NEAA, 55 mM b-mercaptoethanol, 25 mg/mL Insulin, and 2 mg/mL heparin. First, iPSCs were directed toward a neural progenitor fate in base media supplemented with 10 µM SB31542 and 2 µM DMH1, with daily media change over eight days. Followed by patterning with 100 nM RA and 1 µg/mL PMN in base media with daily media change, for four days. Cells were then transferred from monolayer to suspension culture in the same medium for an additional eight days to form neurospheres. Oligodendrocyte precursor cells (OPCs) were then generated and expanded in suspension using OPC media (base media supplemented with 10 ng/mL PDGF-AA, 10 ng/mL IGF-1, 10 ng/mL NT-3, 5 ng/mL HGF, 60 ng/mL 3, 3′, 5-Triiodo-L-thyronine (T3), 100 ng/mL biotin, and 1 µM db-cAMP) for 10 days. Following this, the cells were plated in OPC media onto poly-L-ornithine/laminin-coated vessels with half media changes performed every other day to support further differentiation toward the OPC lineage, which continues until day 75. At this stage, cells were detached and passaged using TrypLE™ Express Enzyme. In the final differentiation step, cells were exposed to ScienCell Astrocyte Medium supplemented with ScienCell Astrocyte Growth Supplements (AGS) for 10 days. AGS was then withdrawn, and final maturation completed by reintroducing IGF-1 and T3 at the previously described concentrations for a further 10 days, with half media changes performed every other day. At this point, oligodendrocytes were considered mature and collected for WB.

#### Astrocyte differentiation

Astrocytes were differentiated from iPSCs following a previously published protocol [43]. First, iPSCs were differentiated into neural progenitor cells following the steps described above for the generation of M-NPCs. M-NPCs were dissociated as single cells and plated at a low cell density (15,000 cells/cm^2^) onto a matrigel coated vessel in M-NPC expansion media (as described above). The next day, medium was replaced with Astrocyte Differentiation Medium (ScienCell Astrocyte Medium containing AGS, 1% fetal bovine serum (FBS) (ScienCell Research Laboratories), 1% penicillin streptomycin (Wisent Bioproducts)). Cells were passaged in the same media at a ratio of 1:2 once per week and were maintained under these culture conditions for 30 days, with a half media change performed every 3-4 days. FBS was then withdrawn to promote astrocyte maturation. Cells were maintained for an additional four weeks with a half media change performed twice per week prior to astrocyte collection for WB.

#### Microglia differentiation

iPSC-derived microglia were differentiated following a published protocol [44]. Briefly, iPSCs were differentiated to hematopoietic precursor cells (iHPCs) using the STEMCELL Technologies STEMdiff™ Hematopoietic Kit. iHPCs were harvested on days 10 and 12 of iHPC differentiation and seeded on matrigel-coated 6-well plates in microglia differentiation media (MDM, DMEM/F12 HEPES no phenol red supplemented with 2X Insulin-transferrin-selenite, 2X B27, 0.5X N2, 1X Glutamax, 1X NEAA, 400 µM monothioglycerol, and 5 µg/mL insulin) + three cytokine cocktail (25 ng/mL M-CSF, 100 ng/mL IL-34, and 50 ng/mL TGF-β1) for further differentiation into microglia. Cells were supplemented with MDM + three cytokine cocktail every other day, with a full media change performed on day 12, for 24 days of microglia differentiation. On day 25 a full media change to MDM + five cytokine cocktail (25 ng/mL M-CSF, 100 ng/mL IL-34, 50 ng/mL TGF-β1, 100 ng/mL CD200, and 100 ng/mL CX3CL1) was performed, followed by supplementation with MDM + five cytokine cocktail on day 27. Microglia were considered mature and utilized for downstream experiments after 28 days of microglial differentiation.

#### Flow cytometry analysis of CD45 and CD11b

Mature iPSC-derived microglia were collected by combining non-adherent cells recovered from culture media with adherent cells gently detached in PBS. Undifferentiated iPSCs were dissociated into single-cell suspensions using Accutase. Cells were first incubated with Live/Dead Fixable Aqua viability dye in PBS for 30 minutes at room temperature, followed by washing in FACS buffer (PBS supplemented with 1% FBS and 0.1% sodium azide). Fc receptor blocking was performed using TrueStain blocking reagent for 10 minutes prior to antibody staining.

Cells were then incubated with fluorophore-conjugated antibodies against CD45 (Alexa Fluor 700; 1:40) and CD11b (PE; 1:20) for 15 minutes at room temperature. After washing, cells were resuspended in FACS buffer and analyzed using a Thermo Attune NxT flow cytometer equipped with 405, 488, and 561 nm lasers (NeuroEDDU Flow Cytometry Facility, McGill University). A minimum of 50,000 events per sample were acquired. Analysis was performed in FlowJo (BD Biosciences), with sequential gating applied to exclude debris and doublets, followed by selection of live cells prior to assessment of CD45 and CD11b expression.

#### Immunofluorescence staining for IBA1 and PU.1

Mature iPSC-derived microglia were plated onto glass coverslips and fixed in 4% paraformaldehyde for 20 minutes at room temperature. Following three washes in PBS, cells were permeabilized with 0.2% Triton X-100 in PBS for 10 minutes and blocked in 5% normal donkey serum (NDS) for 1 hour. Primary antibodies against IBA1 (1:1,000 Synaptic Systems 234 009) and PU.1 (1:500) were diluted in blocking solution and incubated overnight at 4°C. After washing in PBS, cells were incubated for 1 hour at room temperature with corresponding secondary antibodies, together with Hoechst (1 μg/mL). Coverslips were washed, mounted, and imaged using a Leica SP8 confocal microscope.

#### RNA extraction and quantitative RT-PCR

Total RNA was isolated using TRIzol reagent followed by purification with the RNeasy Mini Kit (Qiagen 74106) according to the manufacturer’s instructions. RNA concentration and purity were assessed by spectrophotometry. Reverse transcription was performed using 500 ng total RNA with M-MLV reverse transcriptase, random hexamers, RNase inhibitor, and dNTPs in first-strand buffer supplemented with DTT. Reactions were incubated at 42°C for 60 minutes followed by enzyme inactivation at 75°C for 10 minutes.

Quantitative PCR was performed using TaqMan gene expression assays (ThermoFisher) on a QuantStudio 3 Real-Time PCR System. Relative expression levels were calculated using the comparative Ct (ΔΔCt) method with GAPDH as the reference gene.

The following TaqMan probes were used: GAPDH–Hs02786624_g1; AIF1 (IBA1)–Hs00610419_g1; OLFML3–Hs01113293_g1; SPI1 (PU.1)–Hs02786711_m1; CX3CR1–Hs01922583_s1; MERTK– Hs01031969_m1; TREM2–Hs00219132_m1; CSF1–Hs00174164_m1; P2RY12–Hs01881698_s1; RUNX1–Hs02558380_s1.

## Key resources table

| REAGENT or RESOURCE | SOURCE | IDENTIFIER |
| --- | --- | --- |
| **Antibodies** |  |  |
| HRP-Goat polyclonal anti-Rabbit antibody Secondary Antibody (H+L) | Thermo Fisher Scientific | CAT#65-6120; RRID: AB_2533967 |
| HRP-Goat polyclonal anti-Mouse antibody Secondary Antibody (H+L) | Thermo Fisher Scientific | CAT#62-6520; RRID: AB_2533947 |
| IRDye® 680CW Goat polyclonal anti-Rabbit Secondary Antibody (H+L) | LI-COR Biosciences | CAT#926-68071; RRID:NA |
| IRDye® 680CW Goat polyclonal anti-Mouse Secondary Antibody (H+L) | LI-COR Biosciences | CAT#926-68070; RRID:NA |
| IRDye® 800CW Goat polyclonal anti-Rabbit Secondary Antibody (H+L) | LI-COR Biosciences | CAT#926-32211; RRID:NA |
| IRDye® 800CW Goat polyclonal anti-Mouse Secondary Antibody (H+L) | LI-COR Biosciences | CAT#926-32210; RRID:NA |
| HRP-Goat multiclonal recombinant anti-Rabbit Secondary Antibody (H+L) | Proteintech | CAT#RGAR001; RRID: AB_3073505 |
| HRP-Goat multiclonal recombinant anti-Mouse Secondary Antibody (H+L) | Proteintech | CAT#RGAM001; RRID: AB_3068333 |
| CoraLite Plus 555-Goat multiclonal recombinant Anti-Rabbit Secondary Antibody (H+L) | Proteintech | CAT# RGAR003; RRID: AB_3073507 |
| CoraLite Plus 555-Goat multiclonal recombinant Anti-Mouse Secondary Antibody (H+L) | Proteintech | CAT# RGAM003; RRID: AB_3068539 |
| Rabbit recombinant monoclonal anti-CHMP2B (clone JE54-35) | Thermo Fisher Scientific | CAT#MA5-36184; RRID: AB_2890433 |
| Rabbit recombinant monoclonal anti-FIG4 (clone N202/7R) | Addgene | CAT#192037; RRID: AB_2916241 |
| Mouse monoclonal anti-FUS (clone CL0190) | Abcam | CAT#ab154141; RRID: AB_2885092 |
| Mouse monoclonal anti-HNRNPA1 (Clone 4B10) | Bio-Techne (Novus Biologicals) | CAT#NB100-672; RRID: AB_524308 |
| Rabbit recombinant monoclonal anti-MATR3 (clone EPR10634(B)) | Abcam | CAT#ab151739; RRID: AB_2885091 |
| Rabbit polyclonal anti-NEK1 | GeneTex | CAT#GTX130828; RRID: AB_2886357 |
| Mouse monoclonal anti-OPTN (clone 6C1H4) | Proteintech | CAT#60293-1-Ig; RRID: AB_2881408 |
| Rabbit recombinant monoclonal anti-SIGMAR1 (Clone D4J2E) | Cell Signaling Technology | CAT#61994; RRID: AB_2799617 |
| Rabbit recombinant monoclonal anti-SOD1 (Clone EP1727Y) (Fluorescent WB) | Abcam | CAT#ab51254; RRID: AB_882757 |
| Rabbit polyclonal anti-SOD1 (Chemiluminescent WB) | GeneTex | CAT#GTX100554; RRID: AB_10618670 |
| Rabbit recombinant monoclonal anti-SQSTM1 (clone EPR4844) (Fluorescent WB) | Abcam | CAT#ab109012; RRID: AB 2810880 |
| Mouse monoclonal anti-SQSTM1 (clone 2C11) (Chemiluminescent WB) | Abcam | CAT#ab56416; RRID: AB 945626 |
| Rabbit recombinant monoclonal anti-TAF15 (clone JE61-92) | Thermo Fisher Scientific | CAT#MA5-44822; RRID: AB 2931279 |
| Rabbit recombinant monoclonal anti-TARDBP (clone 11N20) | Proteintech | CAT#80001-1-RR; RRID: AB 2882933 |
| Rabbit recombinant monoclonal anti-TBK1 (Clone EP611Y) (Chemiluminescent WB) | Abcam | CAT#ab40676; RRID: AB 776632 |
| Rabbit recombinant monoclonal anti-TBK1 (clone E8I3G) ((Fluorescent WB) | Cell Signaling Technology | CAT#38066; RRID: AB 2827657 |
| Rabbit recombinant monoclonal anti-TIA1 (Clone D1Q3K) | Cell Signaling Technology | CAT#86050; RRID: AB 2800070 |
| Mouse monoclonal anti-UBQLN2 (clone 6H9) | Abcam | CAT#ab190283; RRID: AB 2747782 |
| Rabbit recombinant monoclonal anti-VCP (Clone 7F3) (Chemiluminescent WB) | Cell Signaling Technology | CAT#2649; RRID: AB 2214629 |
| Rabbit recombinant monoclonal anti-VCP (clone EPR3307(2)) (Fluorescent WB) | Abcam | CAT#ab109240; RRID: AB 10862588 |
| Rabbit polyclonal anti-ANXA11 | Aviva Systems Biology | CAT#ARP87950_P050; RRID: AB 2894925 |
| Mouse monoclonal anti-C9orf72 (clone GT1553) | GeneTex | CAT#GTX634482; RRID: AB 2784545 |
| Rabbit polyclonal anti-CHCHD10 (Chemiluminescent WB) | Proteintech | CAT#25671-1-AP; RRID: AB 2880187 |
| Mouse monoclonal anti-CHCHD10 (clone OTI3B8) (Fluorescent WB) | Thermo Fisher Scientific | CAT#MA5-27535; RRID: AB 2724132 |
| Rabbit recombinant monoclonal anti-LGALS1 (clone EPR3206(2)) | Abcam | CAT#ab138513; RRID: AB 2894851 |
| Rabbit recombinant monoclonal anti-PFN1 (clone ARC1469) | ABclonal | CAT#A9188; RRID: AB 2863682 |
| Rabbit recombinant monoclonal anti-SPG11 (clone 2H25L11) | Thermo Fisher Scientific | CAT#702776; RRID: AB 2725406 |
| Mouse monoclonal anti-ACSL5 (Clone 5H8) | Abnova | CAT#H00051703-M01; RRID: AB 509241 |
| Rabbit recombinant monoclonal anti-CAV1 (clone SZ02-01) (Chemiluminescent WB) | Thermo Fisher Scientific | CAT#MA5-32020; RRID: AB 2809314 |
| Rabbit recombinant monoclonal anti-CAV1 (Clone ARC50848) (Fluorescent WB) | ABclonal | CAT#A19006; RRID: AB 2862498 |
| Rabbit recombinant monoclonal anti-ATXN2 (clone EPR23630-49) | Abcam | CAT#ab254362; RRID: AB 2885130 |
| Rabbit recombinant monoclonal anti-TUBA4A (Clone EPR13478(B)) (Chemiluminescent WB) | Abcam | CAT#ab176560; RRID: AB 2860019 |
| Rabbit recombinant monoclonal anti-TUBA4A (Clone JM73-24) (Fluorescent WB) | Thermo Fisher Scientific | CAT#MA5-32738; RRID: AB 2810015 |
| Rabbit recombinant monoclonal anti-VAPB (clone EPR27026-50) (Chemiluminescent WB) | Abcam | CAT#ab315013; RRID: AB 3086770 |
| Rabbit polyclonal anti-VAPB (Fluorescent WB) | Thermo Fisher Scientific | CAT#PA5-53023; RRID: AB 2649391 |
| Rabbit recombinant monoclonal anti-ALS2 (clone EPR11185) | Abcam | CAT#ab170896; RRID: AB 2885099 |
| Mouse monoclonal anti-HNRNPA2B1 (clone DP3B3) | Bio-Techne (Novus Biologicals) | CAT#NB120-6102; RRID: AB 790226 |
| Rabbit recombinant monoclonal anti-KIF5A (clone 241124H1) | Proteintech | CAT#84013-4-RR; RRID: AB 3671581 |
| Rabbit recombinant monoclonal anti-Synaptotagmin 1 (clone HL1626) | GeneTex | CAT#GTX637119; RRID: AB 3068350 |
| Rabbit recombinant monoclonal anti-CD44 (clone EPR18668) | Abcam | CAT# ab189524; RRID: AB 2885107 |
| Rabbit recombinant monoclonal anti-ISL1 (Clone EPR10362) | Abcam | CAT#ab178400; RRID: AB 2927537 |
| Rat recombinant monoclonal anti-Myelin Basic Protein antibody (clone 12) | Abcam | CAT#ab7349; RRID: AB 305869 |
| Rabbit polyclonal anti-Tyrosine Hydroxylase | Pel-Freeze | CAT#P40101-150; RRID:<br>AB_2617184 |
| Rabbit recombinant monoclonal anti-CD11b (clone EPR1344) | Abcam | CAT#ab133357; RRID:<br>AB_2650514 |
| Rabbit recombinant monoclonal anti-IBA1 (clone E4O4W) (WB) | Cell Signaling Technology | CAT#17198; RRID:<br>AB_2820254 |
| Chicken monoclonal recombinant anti-IBA1 (clone Ch311H9) (IF) | Synaptic Systems | CAT#234 009<br>AB_2891282 |
| Mouse monoclonal anti-CD11b, pre-conjugated with PE (clone ICRF44) | BD Biosciences | CAT#555388<br>AB_395789 |
| Mouse monoclonal anti-CD45, pre-conjugated with Alexa Fluor 700 (clone HI30) | BioLegend | CAT#304024<br>AB_493761 |
| Rabbit polyclonal anti-PU.1 | Cell Signaling Technology | CAT#2266<br>AB_10692379 |
| **Other reagents** |  |  |
| Accutase | STEMCELL Technologies | CAT#07922 |
| Astrocyte Growth Supplements (AGS) | ScienCell Research Laboratories | CAT#1852 |
| Astrocyte Medium (AM) | ScienCell Research Laboratories | CAT#1801-b |
| BLUelf Prestained Protein Ladder | FroggaBio | CAT#PM008 |
| CellTracker deep red dye | Thermo Fisher Scientific | CAT#C34565 |
| CellTracker green chloromethyl derivative of fluorescein diacetate (CMFDA) dye | Thermo Fisher Scientific | CAT#C2925 |
| Centrifugal filter unit, Amicon Ultra-15 | MilliporeSigma | CAT#UFC901096 |
| Live/Dead Fixable Aqua viability dye | Thermo Fisher Scientific | CAT#L34965 |
| mTeSR1 | STEMCELL Technologies | CAT#85850 |
| NuPAGE™ Bis-Tris Midi Protein Gels, 10%, 1.0 mm | Thermo Fisher Scientific | CAT#WG1201BOX |
| NuPAGE™ MES SDS Running Buffer (20X) | Thermo Fisher Scientific | CAT#NP000202 |
| NuPAGE™ MOPS SDS Running Buffer (20X) | Thermo Fisher Scientific | CAT#NP000102 |
| PhenoPlate, 96-well, optically clear flat bottom | Perkin Elmer | CAT#6055300 |
| Pierce™ IP Lysis Buffer | Thermo Fisher Scientific | CAT#87788 |
| RIPA Lysis and Extraction Buffer | Thermo Fisher Scientific | CAT#89901 |
| RNeasy Mini Kit | Qiagen | CAT#74106 |
| STEMdiff™ Forebrain Neuron Differentiation Kit | STEMCELL Technologies | CAT#08600 |
| STEMdiff™ Forebrain Neuron Maturation Kit | STEMCELL Technologies | CAT#08605 |
| STEMdiff™ Hematopoietic Kit | STEMCELL Technologies | CAT#05310 |
| STEMdiff™ Hematopoietic Kit | STEMCELL Technologies | CAT#05310 |
| STEMdiff™ Hematopoietic Kit | STEMCELL Technologies | CAT#05310 |
| TRIzol | Thermo Fisher Scientific | CAT#15596026 |
| TrueStain blocking reagent | BioLegend | CAT#422302 |
| **Experimental models: Cell lines** |  |  |
| Healthy control iPSCs from the AIW002-02 background | EDDU, McGill University | NA |
| Healthy control iPSCs from the QPN929 background | EDDU, McGill University | NA |
| **Software and algorithms** |  |  |
| LI-COR Image Studio Lite | licorbio.com/image-studio | NA |
| FIJI Software | fiji.sc/ | RRID:SCR_002285 |
| MetaXpress High-Content Screening Software version 6.7.1.157 | moleculardevices.com/products/cellular-imaging-systems/high-content-analysis/metaxpress-overview | RRID:SCR_016654 |
| PyCharm 2023.3.4 | jetbrains.com/pycharm/ | RRID:SCR_018221 |
| RStudio 2023.09.1 | cran.rstudio.com/ | RRID:SCR_000432 |

| Other |  |  |
| --- | --- | --- |
| Antibody Registry | antibodyregistry.org | RRID:SCR_006397 |
| Cellosaurus - a knowledge resource on cell lines | cellosaurus.org | RRID:SCR_013869 |
| CiteAb - Reagent Search & Life Science Data | citeab.com | RRID:SCR_009653 |
| ClinGen | https://clinicalgenome.org/ | RRID:SCR_014968 |
| DepMap: The Cancer Dependency Map Project at Broad Institute | depmap.org/portal | RRID:SCR_017655 |
| PubMed | pubmed.ncbi.nlm.nih.gov | RRID:SCR_004846 |
| Structural Genomics Consortium | https://www.thesgc.org/ | RRID:SCR_003890 |
| The Human Protein Atlas | proteinaatlas.org | RRID:SCR_006710 |
| Zenodo | https://zenodo.org/ | RRID:SCR_004129 |

## Resource availability

- All 33 ALS-RAP antibody characterization reports are publicly available on Zenodo, together with the complete raw characterization datasets (WB, IP, and IF, including Galectin-1): https://zenodo.org/communities/ycharos/
- A subset of 15 ALS-RAP targets has also been published and peer-reviewed in the *F1000Research* YCharOS Gateway and compiled within the dedicated ALS area: https://f1000research.com/gateways/ycharos/als
- Raw WB data supporting ALS-RAP protein expression profiling across neurological cell types (Figure 4) are available on Zenodo.

`o` The chemiluminescent-based experiment: https://doi.org/10.5281/zenodo.18484385
`o` The fluorescent-based experiment: https://doi.org/10.5281/zenodo.18484483
- Curated YCharOS datasets, together with application-specific antibody recommendations, are accessible through the Only Good Antibodies (OGA) website: https://onlygoodantibodies.co.uk/

## Acknowledgements

We would like to thank Kathleen Southern for the coordination of information exchange between laboratories involved in the ALS-RAP project. We would like to thank Abcam, ABCD Antibodies, ABclonal, Aviva Systems Biology, Cell Signaling Technology, Developmental Studies Hybridoma Bank, GeneTex, Revvity, Institute for Protein Innovation, MilliporeSigma, Novus Biologicals, Proteintech, R&D Systems, Thermo Fisher Scientific for providing antibodies and/or KO cell lines, free of charge, for ALS-RAP related targets. We would like to acknowledge the Birth Defects Research Laboratory, University of Washington, Seattle, USA for providing fetal tissues.

This work was supported by a grant from the Motor Neurone Disease Association, the ALS Association and ALS Canada to develop the ALS-Reproducibility Antibody Platform. This work was also supported by the Government of Canada through Genome Canada, Genome Quebec and Ontario Genomics (OGI-210) and by grant from the Quebec Consortium for Drug Discovery (CQDM), a grant from the Ministère de l’Économie, de l’Innovation et de l’Énergie du Québec. This work was supported by a grant from the National Centre for the Replacement, Refinement and Reduction of Animals in Research (NC3Rs) and MRC (NC3Rs Ref: NC/NAM0019/1, MRC UKRI076) alongside support from the Leicester Institute for Precision Health. The Structural Genomics Consortium is a registered charity (no. 1097737) that receives funds from Bayer, Boehringer Ingelheim, Bristol Myers Squibb, Genentech, Genome Canada through Ontario Genomics Institute (grant no. OGI-196), the European Union (EU) and European Federation of Pharmaceutical Industries and Associations through the Innovative Medicines Initiative 2 Joint Undertaking (EUbOPEN grant no. 875510), Janssen, Merck (also known as EMD in Canada and the USA), Pfizer and Takeda. R.A. is supported by a Mitacs postdoctoral fellowship. This work was supported by a CIHR project grant (PJT-195804). E.A.F. is supported by a Canada Research Chair (Tier 1) in Parkinson’s disease. J.P.R. is supported by an ALS Scholars in Therapeutics postdoctoral fellowship from the Sean M. Healey & AMG Center for ALS at Massachusetts General Hospital (MGH), ALS FindingACure, and FightMND.

## Author contribution

R.A., O.G., A.M.E., T.M.D., P.S.M., and C.L. conceived the study. I.M. analyzed ALS-RAP RNA sequencing data from the Human Protein Atlas. R.A., C.Z., S.G.B., V.R.M., C.A., and M.F. performed experiments and generated data. E.J.M., M.C., C.X.-Q.C., V.E.C.P., and V.S. generated iPSC-derived neurological cell types. M.S.G, B.T.F. and H.S.V established the Only Good Antibodies (OGA) website. T.K., W.H.L., E.W., and C.A.M. generated and characterized scFv antibodies. B.D.M. and L.K. established curated the ALS-RAP target list. M.-F.D. and L.M.H. isolated fetal astrocytes and fetal microglia and generated monocyte-derived macrophages. V.F., R.A., S.G.B and C.L. participated in data dissemination. P.A.D., J.P.R. and G.A.R. contributed to defining the initial ALS-RAP target list. R.A., I.M., and C.L. analyzed and interpreted the data. C.L., O.G., S.G., A.M.E., T.M.D., E.A.F., W.H.L. and P.S.M. provided resources and supervision. C.L. wrote the original draft of the manuscript with input from R.A., A.M.E. and P.S.M. All authors reviewed and approved the final manuscript.

## Declaration of interests

The authors report no competing interests.

## Supplemental information

- Supplementary Figures 1-5
- Supplementary Table 1 (Table S1). List of 303 antibodies targeting ALS-RAP proteins evaluated in this study, together with the DOI linking to the corresponding antibody characterization reports.
- Supplementary Table 2 (Table S2). Sequences of synthetic ScFv antibodies

---

**Figure S1:**
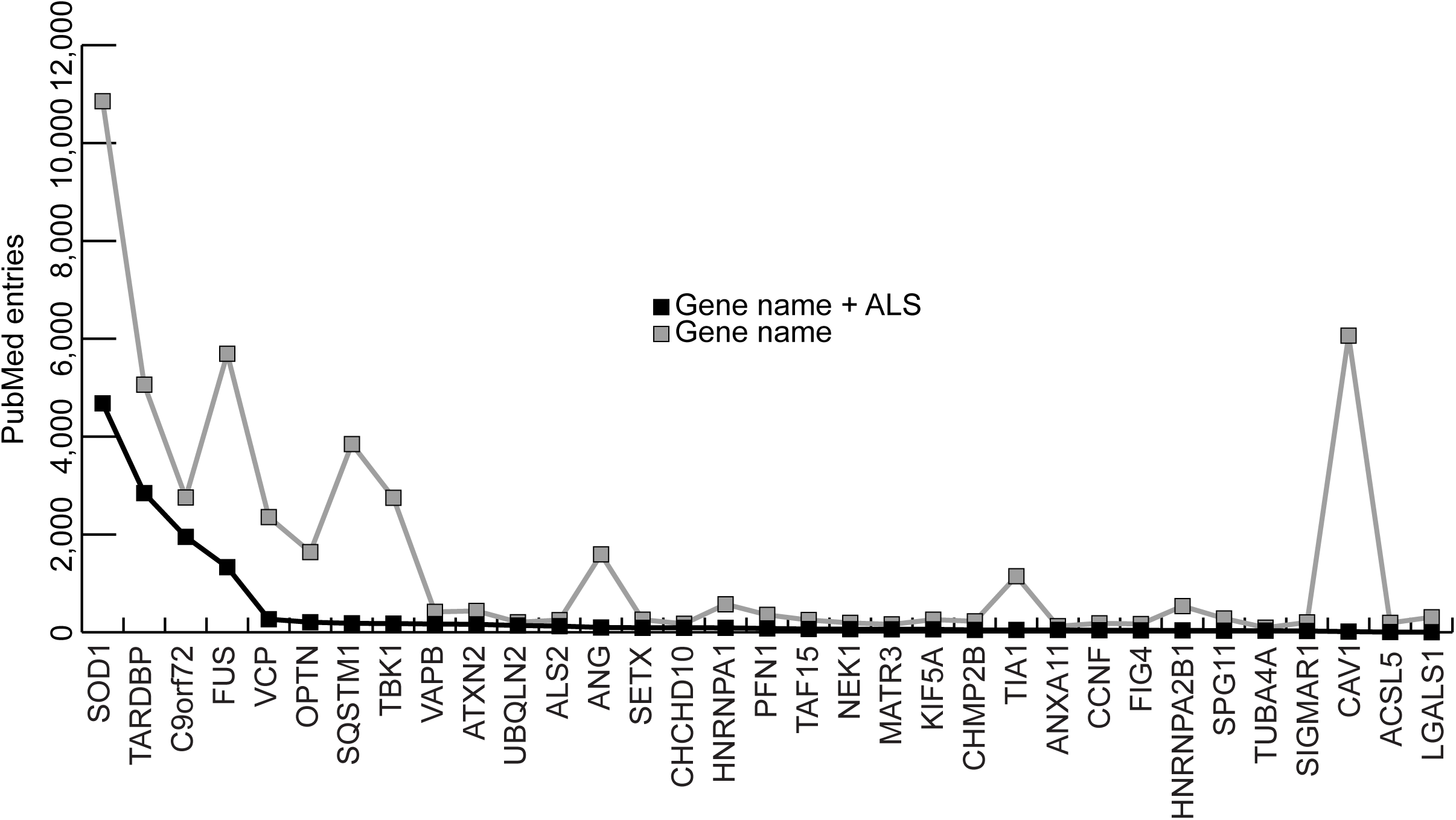
Characterization of ALS-RAP targets. Number of PubMed entries related to ALS-RAP genes. Black points represent the number of entries retrieved using the search term “gene name + ALS,” while gray points represent entries retrieved using the gene name alone. The x-axis indicates the number of PubMed entries, and the y-axis lists the gene names.

**Figure S2:**
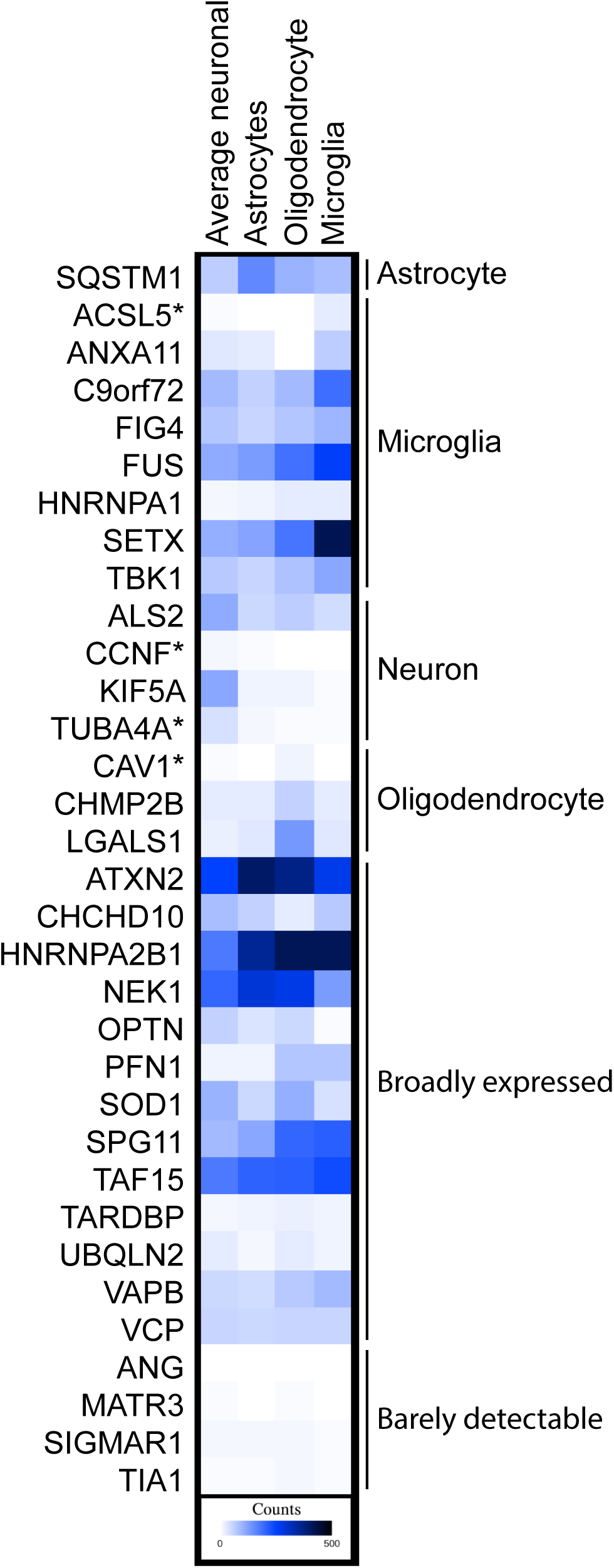
ALS-RAP RNA expression across brain cell types. Heatmap showing normalized RNA expression values from the Human Protein Atlas (HPA) single nuclei brain dataset. Expression is given as nTPM values on a scale from 0-500. Genes denoted with an asterisk (*) are classified as “barely detectable” but show substantial expression in a single cell type.

**Figure S3:**
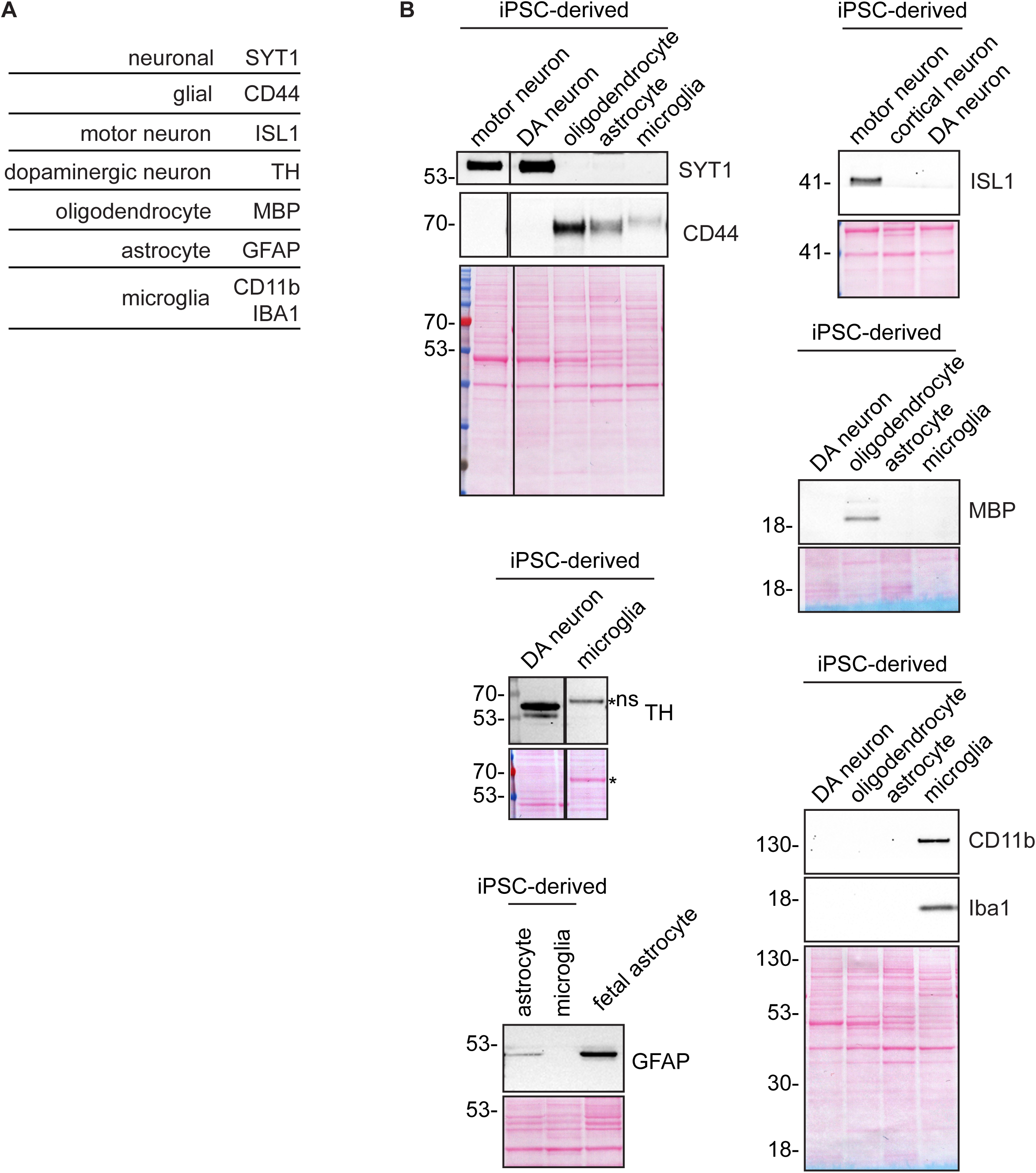
Protein-level characterization of iPSC-derived neurological cells. **(A)** Table listing cell type–specific marker proteins. **(B)** WB analysis of marker proteins using antibodies against the indicated proteins. A Ponceau-stained membrane is shown to assess protein loading and transfer efficiency. Lysates were prepared from the indicated human iPSC-derived cell types or fetal astrocyte culture. Vertical lines between the motor neuron and DA neuron lanes and between the DA neuron and microglia lanes indicate that WB were spliced to remove additional conditions not relevant to this study. DA, dopaminergic; NPC, neuronal progenitor cells.

**Figure S4:**
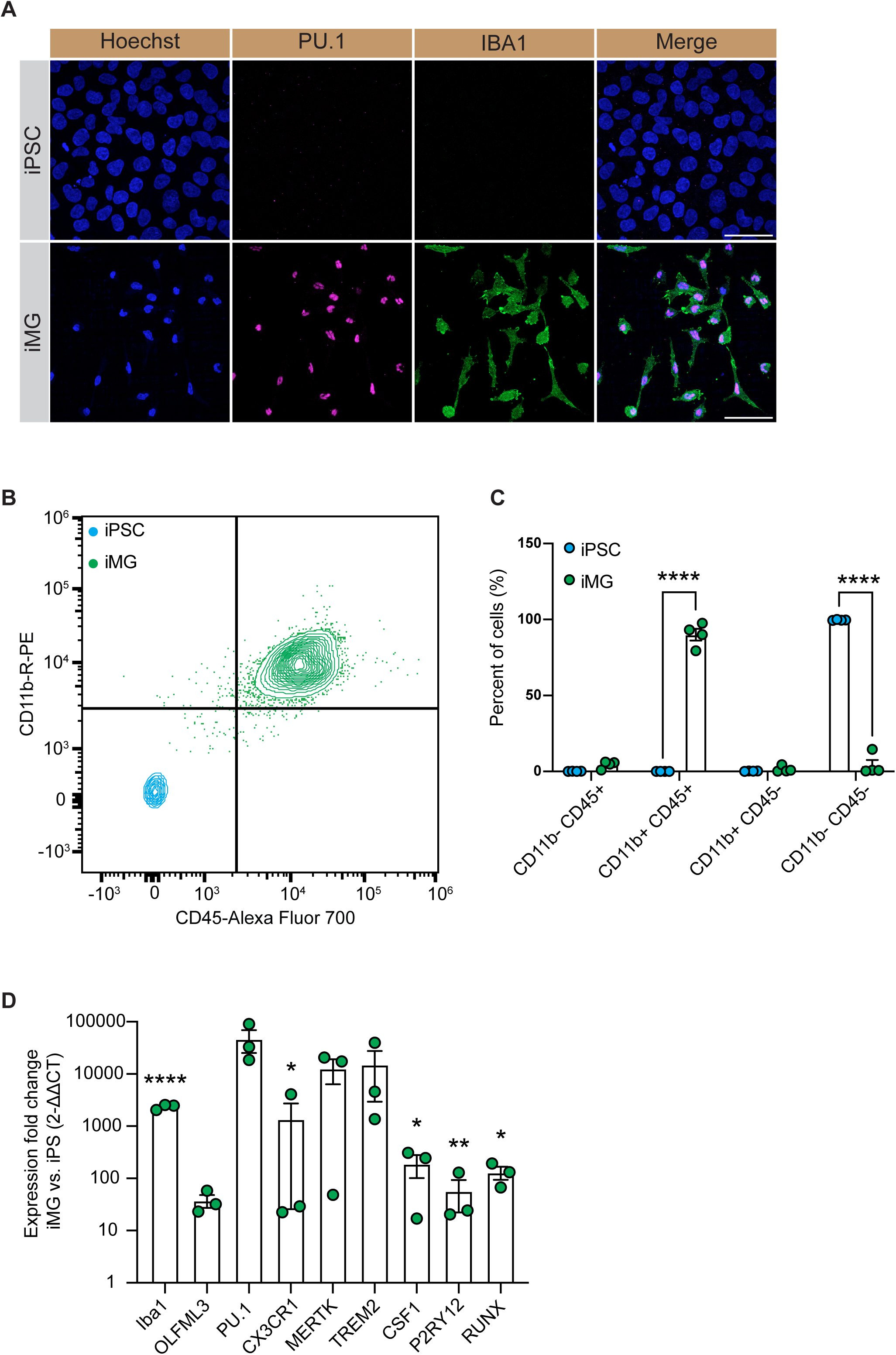
Quality control and marker validation of iPSC-derived microglia. **(A)** Immunofluorescence staining showing expression of IBA1 and PU.1 in iMGs, with no detectable signal in undifferentiated iPSCs. **(B)** Representative flow cytometry plots demonstrating expression of CD45 and CD11b in iMGs but not in iPSCs. **(C)** Quantification of the percentage of CD45-positive, CD11b-positive, and double-positive cells in iPSCs and iMGs, as measured by flow cytometry (N = 4). Statistical analysis was performed using two-way ANOVA with Bonferroni post hoc test. **(D)** RT-qPCR expression levels of canonical microglial markers in iMGs compared to iPSCs (N = 3). Statistical analysis was performed using Student’s t-test. iMG, iPSC-derived microglia; iPSC, induced pluripotent stem cell. * p ≤ 0.05, ** p ≤ 0.01, *** p ≤ 0.001, **** p ≤ 0.0001. Scale bar, 50 µm

**Figure S5:**
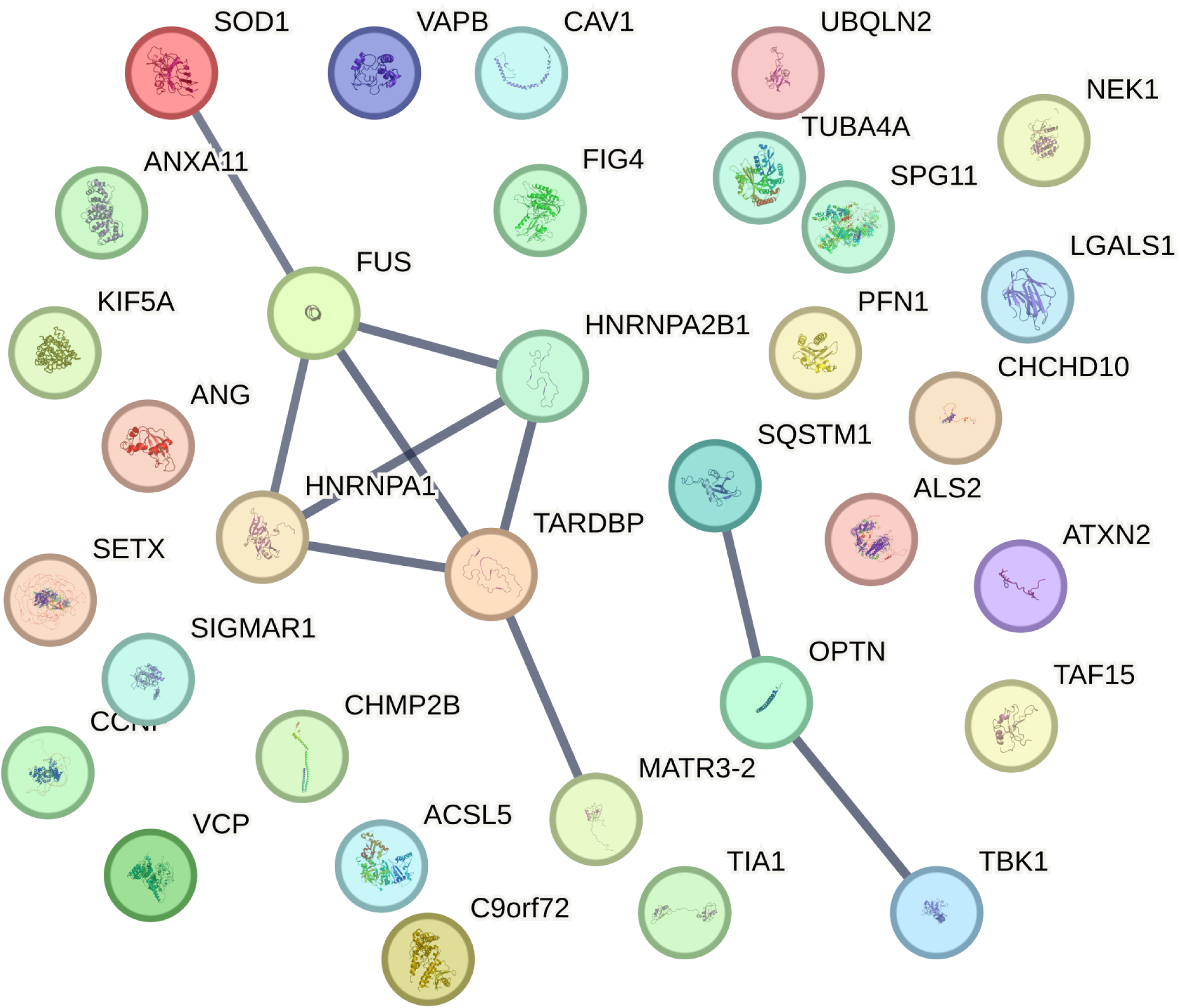
STRING analysis of known physical interactions between ALS-RAP proteins.

## Notes

### Competing Interest Statement

The authors have declared no competing interest.

### Summary of Updates

This version includes revisions made in response to eLife peer review and corresponds to the eLife Version of Record.

## References

1. Kahn, R.A., H.S. Virk, and P.S. McPherson, Heed a decade of calls for antibody validation. Nature, 2023. 620(7974): p. 492.

2. Biddle, M., et al., Improving the integrity and reproducibility of research that uses antibodies: a technical, data sharing, behavioral and policy challenge. MAbs, 2024. 16(1): p. 2323706.

3. Baker, M., Reproducibility crisis: Blame it on the antibodies. Nature, 2015. 521(7552): p. 274–6.

4. Goodman, S.L., The antibody horror show: an introductory guide for the perplexed. N Biotechnol, 2018. 45: p. 9–13.

5. Laflamme, C., et al., Implementation of an antibody characterization procedure and application to the major ALS/FTD disease gene C9ORF72. Elife, 2019. 8.

6. Ayoubi, R., et al., Scaling of an antibody validation procedure enables quantification of antibody performance in major research applications. Elife, 2023. 12.

7. Nijs, M. and P. Van Damme, The genetics of amyotrophic lateral sclerosis. Curr Opin Neurol, 2024. 37(5): p. 560–569.

8. Rehm, H.L., et al., ClinGen--the Clinical Genome Resource. N Engl J Med, 2015. 372(23): p. 2235–42.

9. Ayoubi, R., et al., A consensus platform for antibody characterization. Nat Protoc, 2024.

10. Monteiro, F.L., J.L.A. Voskuil, and C. Williams, YCharOS protocol for antibody validation. Nat Protoc, 2025. 20(6): p. 1389–1390.

11. Lund-Johansen, F., A strong case for third-party testing. Elife, 2023. 12.

12. Laflamme, C., et al., Opinion: Independent third-party entities as a model for validation of commercial antibodies. N Biotechnol, 2021. 65: p. 1–8.

13. Hornsby, M., et al., A High Through-put Platform for Recombinant Antibodies to Folded Proteins. Mol Cell Proteomics, 2015. 14(10): p. 2833–47.

14. Marx, V., Change-makers bring on recombinant antibodies. Nat Methods, 2020. 17(8): p. 763–766.

15. Preger, C., et al., Generation and validation of recombinant antibodies to study human aminoacyl-tRNA synthetases. J Biol Chem, 2020. 295(41): p. 13981–13993.

16. Marcon, E., et al., Assessment of a method to characterize antibody selectivity and specificity for use in immunoprecipitation. Nat Methods, 2015. 12(8): p. 725–31.

17. Ayoubi, R., et al., The identification of high-performing antibodies for Profilin-1 for use in Western blot, immunoprecipitation and immunofluorescence. F1000Res, 2023. 12: p. 348.

18. Biddle, M.S. and H.S. Virk, YCharOS open antibody characterisation data: Lessons learned and progress made. F1000Res, 2023. 12: p. 1344.

19. Bandrowski, A., et al., The Antibody Registry: ten years of registering antibodies. Nucleic Acids Res, 2023. 51(D1): p. D358–D367.

20. Bairoch, A., The Cellosaurus, a Cell-Line Knowledge Resource. J Biomol Tech, 2018. 29(2): p. 25–38.

21. Cai, Z., et al., Association between abnormal expression and methylation of LGALS1 in amyotrophic lateral sclerosis. Brain Res, 2022. 1792: p. 148022.

22. Kato, T., et al., Galectin-1 is a component of neurofilamentous lesions in sporadic and familial amyotrophic lateral sclerosis. Biochem Biophys Res Commun, 2001. 282(1): p. 166–72.

23. Karlsson, M., et al., A single-cell type transcriptomics map of human tissues. Sci Adv, 2021. 7(31).

24. McKeever, P.M., et al., Single-nucleus transcriptome atlas of orbitofrontal cortex in ALS with a deep learning-based decoding of alternative polyadenylation mechanisms. Cell Genom, 2025: p. 101007.

25. Gygi, S.P., et al., Correlation between protein and mRNA abundance in yeast. Mol Cell Biol, 1999. 19(3): p. 1720–30.

26. Vogel, C. and E.M. Marcotte, Insights into the regulation of protein abundance from proteomic and transcriptomic analyses. Nat Rev Genet, 2012. 13(4): p. 227–32.

27. Bellenguez, C., et al., New insights into the genetic etiology of Alzheimer’s disease and related dementias. Nat Genet, 2022. 54(4): p. 412–436.

28. Zhang, J., et al., Genome-wide association study in Alzheimer’s disease: a bibliometric and visualization analysis. Front Aging Neurosci, 2023. 15: p. 1290657.

29. Leonard, H.L. and P. Global Parkinson’s Genetics, Novel Parkinson’s Disease Genetic Risk Factors Within and Across European Populations. medRxiv, 2025.

30. Kim, J.J., et al., Multi-ancestry genome-wide association meta-analysis of Parkinson’s disease. Nat Genet, 2024. 56(1): p. 27–36.

31. van Rheenen, W., et al., Common and rare variant association analyses in amyotrophic lateral sclerosis identify 15 risk loci with distinct genetic architectures and neuron-specific biology. Nat Genet, 2021. 53(12): p. 1636–1648.

32. Edwards, A.M., et al., Too many roads not taken. Nature, 2011. 470(7333): p. 163–5.

33. Carter, A.J., et al., Target 2035: probing the human proteome. Drug Discov Today, 2019. 24(11): p. 2111–2115.

34. O’Rourke, J.G., et al., C9orf72 is required for proper macrophage and microglial function in mice. Science, 2016. 351(6279): p. 1324–9.

35. Masrori, P., et al., C9orf72 hexanucleotide repeat expansions impair microglial response in ALS. Nat Neurosci, 2025. 28(11): p. 2217–2230.

36. Michaelis, T., et al., Autoimmune response to C9orf72 protein in amyotrophic lateral sclerosis. Nature, 2025.

37. Siletti, K., et al., Transcriptomic diversity of cell types across the adult human brain. Science, 2023. 382(6667): p. eadd7046.

38. Deneault, E., et al., A streamlined CRISPR workflow to introduce mutations and generate isogenic iPSCs for modeling amyotrophic lateral sclerosis. Methods, 2022. 203: p. 297–310.

39. Lepine, S., et al., Homozygous ALS-linked mutations in TARDBP/TDP-43 lead to hypoactivity and synaptic abnormalities in human iPSC-derived motor neurons. iScience, 2024. 27(3): p. 109166.

40. Kriks, S., et al., Dopamine neurons derived from human ES cells efficiently engraft in animal models of Parkinson’s disease. Nature, 2011. 480(7378): p. 547–51.

41. C, X.Q.C., et al., Generation of patient-derived pluripotent stem cell-lines and CRISPR modified isogenic controls with mutations in the Parkinson’s associated GBA gene. Stem Cell Res, 2022. 64: p. 102919.

42. Piscopo, V.E.C., et al., The use of a SOX10 reporter toward ameliorating oligodendrocyte lineage differentiation from human induced pluripotent stem cells. Glia, 2024. 72(6): p. 1165–1182.

43. Soubannier, V., et al., Rapid Generation of Ventral Spinal Cord-like Astrocytes from Human iPSCs for Modeling Non-Cell Autonomous Mechanisms of Lower Motor Neuron Disease. Cells, 2022. 11(3).

44. McQuade, A., et al., Development and validation of a simplified method to generate human microglia from pluripotent stem cells. Mol Neurodegener, 2018. 13(1): p. 67.

